# ISL1 is required for the terminal differentiation and maintenance of Starburst Amacrine Cells

**DOI:** 10.64898/2026.10.05.756830

**Authors:** Yessica C. Santana Agreda, Kevin M. Wright

## Abstract

During mammalian nervous system development, multipotent progenitor cells differentiate into hundreds of distinct neuronal subtypes. Neuronal subtype terminal differentiation is regulated by transcription factors that directly control the acquisition and maintenance of terminal identity. Using the mouse retina as a model system, we show that the transcription factor ISL1 is required for the terminal differentiation and maintenance of starburst amacrine cells (SACs), the sole cholinergic neuron subtype in the retina. We conditionally deleted *Isl1* selectively from amacrine and horizontal cell precursors, allowing us to assess its cell-autonomous role in SAC development without disrupting its critical function in bipolar cell and retinal ganglion cell development. We find that ISL1 is not required for initial SAC specification but is required for their terminal differentiation. *Isl1*-deficient SACs retain a GABAergic amacrine cell identity but fail to adopt many SAC-specific features including expression of genes required for cholinergic transmission. In the absence of ISL1, SACs fail to migrate properly, with both OFF- and ON-SACs localizing to the inner nuclear layer. Deletion of *Isl1* from post-migratory SACs at later stages of development revealed that it is also required for maintaining SAC identity and proper dendrite stratification. Loss of *Isl1* from SACs ultimately caused abnormal development of the direction selective circuit. Together, these findings establish that ISL1 is required for the induction and maintenance of SAC-specific identity from a generic amacrine state.

## INTRODUCTION

Proper neuronal circuit function is dependent on the differentiation and wiring of diverse neuronal subtypes. While many of the temporal and spatial transcriptional pathways that give rise to broad neuronal classes in mammals have been identified, our understanding of the transcription factors that subsequently define the terminal features of individual neuronal subtypes remains incomplete. In the relatively simple nervous system of *C. elegans*, terminal selector transcription factors direct the acquisition and maintenance of neuronal subtype identity via direct *cis*-regulation of effector genes (Hobert, 2016; Leyva-Díaz & Hobert, 2019). These include genes that define a neuronal subtype’s identity and function such as ion channels, neurotransmitter synthesizing enzymes, neurotransmitter receptors, neuropeptides, and synaptic adhesion molecules. In contrast, the extreme level of neuronal subtype diversity and genetic complexity in mammals has hampered our understanding of the role specific transcription factors play in neuronal subtype terminal differentiation (Bakshi et al., 2025; Flames & Hobert, 2011).

The mammalian retina is an excellent model system for investigating neuronal subtype development. It is composed of five main neuronal classes organized in a highly stereotyped laminar structure (Masland, 2001). Photoreceptors (PRs) in the outer retina convert photons into electrical signals that are transmitted to bipolar cells (BCs), which in turn convey information to retinal ganglion cells (RGCs) in the inner retina.

Horizontal cells (HCs) provide inhibition at synapses between PRs and BCs in the outer plexiform layer (OPL) and amacrine cells (ACs) provide inhibition between BCs and RGCs in the inner plexiform layer (IPL) **(Fig. 1A**). In the mouse, these five classes can be further subdivided into ∼130 transcriptionally distinct neuronal subtypes (J. Li et al., 2024; Shekhar et al., 2016). ACs and RGCs show the highest level of transcriptional diversity, with an estimated 63 and 46 putative subtypes, respectively (Rheaume et al., 2018; Tran et al., 2019; Yan et al., 2020). Many of these subtypes are conserved in other mammals (Peng et al., 2019; Tommasini et al., 2026). While the early transcriptional pathways that regulate the development of the five retinal neuron classes are relatively well understood, how specific neuronal subtypes differentiate from their precursors remains less clear (Bassett & Wallace, 2012; Kaplan et al., 2025; S. Liu et al., 2020; Mu et al., 2008).

**Figure 1.**
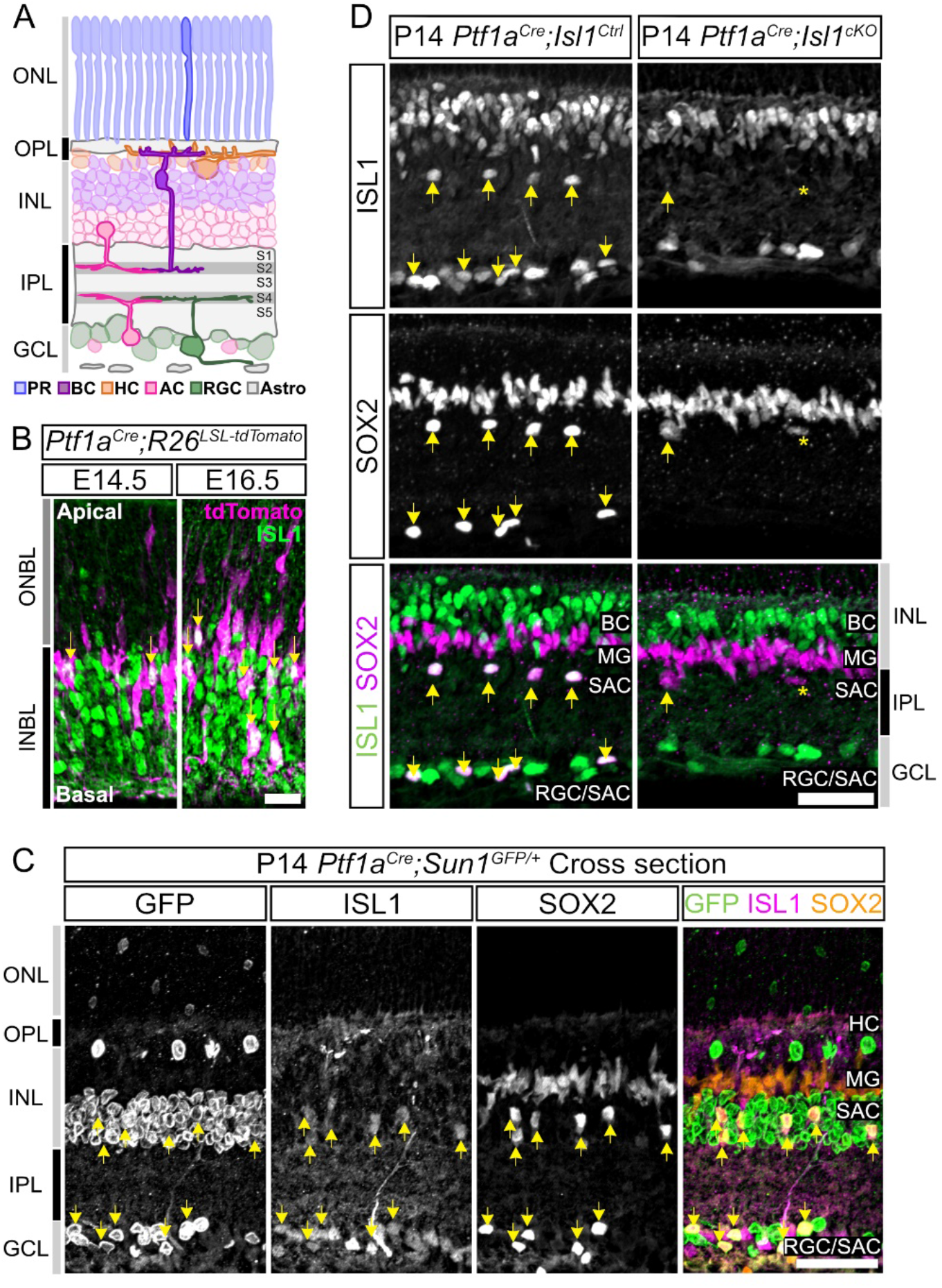
Selective deletion of *Isl1* from SACs using *Ptf1a^Cre^*. (**A**) Schematic of the vertebrate retina detailing the five major neuronal classes and non-neuronal cells. (**B**) Onset of ISL1 in *Ptf1a^Cre^;R26^LSL-tdTomato^* embryonic retinas at E14.5 (n = 5) and 16.5 (n = 3). Yellow arrows indicate SACs (tdT^+^/ISL1^+^). Scale bar = 20µm. (**C**) P14 *Ptf1a^Cre^;Sun1^GFP/+^* cross sections (n = 3) with GFP, ISL1, and SOX2. Yellow arrows indicate SACs (GFP^+^/ISL1^+^/SOX2^+^). Scale bar = 50 μm. (**D**) P14 *Ptf1a^Cre^;Isl1^Ctrl^* (n = 3) and P14 *Ptf1a^Cre^;Isl1^cKO^* (n = 3) retinas immunostained for ISL1 and SOX2. Yellow arrows indicate SACs (ISL1^+^/SOX2^+^) in control (left) and mutant SOX2^+^ SAC (right). Asterisk indicates a SOX2^+^ INL retinal astrocyte. Scale bar = 35 μm. **AC** = amacrine cells, **Astro** = retinal astrocytes, **BC** = bipolar cells, **GCL** = ganglion cell layer, **HC** = horizontal cells, **INBL** = inner neuroblast layer, **INL** = inner nuclear layer, **IPL** = inner plexiform layer, **MG** = Müller glia, **ONBL** = outer neuroblast layer, **ONL** = outer nuclear layer, **OPL** = outer plexiform layer, **PR** = photoreceptors, **RGC** = retinal ganglion cell, **SAC** = starburst amacrine cell, **S1**-**S5** = Sublamina 1-5.

*Isl1* is a widely expressed LIM-homeobox transcription factor important for the development of various neuronal and non-neuronal cell populations (N. Kim et al., 2015). ISL1 is present in all cholinergic neuron populations in the mouse. It is required for the differentiation of cholinergic neurons in the spinal cord and forebrain, and directly regulates a battery of genes required for acetylcholine synthesis and packaging (Cho et al., 2014; Elshatory & Gan, 2008; Liang et al., 2011; Pfaff et al., 1996). The sole cholinergic neuron population in the retina, starburst amacrine cells (SACs), supply both GABAergic and cholinergic inputs to the direction selective (DS) circuit (Elshatory, Deng, et al., 2007; Reed et al., 2026). SACs can be split into two subtypes: OFF-SACs that reside in the inner nuclear layer (INL) and stratify in sublamina 2 (S2) of the IPL, and ON-SACs that reside in the ganglion cell layer (GCL) and stratify in S4 (**Fig. 1A**).

While pan-retinal deletion of *Isl1* suggests that it is required for the development of SACs, it is also required for the survival of the majority of BCs and RGCs (Elshatory, Everhart, et al., 2007). Therefore, the cell-autonomous function of ISL1 in regulating the differentiation of SACs is obscured by the widespread loss of their pre- and post-synaptic partners when *Isl1* is deleted throughout the retina.

Here, we developed a genetic strategy to selectively delete *Isl1* in ACs to minimize off-target effects, allowing us to test the hypothesis that ISL1 functions cell-autonomously to regulate the terminal differentiation of SACs. We find that in the absence of *Isl1*, presumptive SACs are specified and express general GABAergic AC markers but fail to terminally differentiate and no longer express many canonical SAC markers. *Isl1*-deficient SACs fail to migrate correctly, with all SACs localizing to the INL, and a significant proportion are subsequently lost through BAX-mediated apoptosis. We also show that ISL1 is also required in post-migratory SACs to maintain their subtype identity. Ultimately, the failure of SACs to properly differentiate in the absence of *Isl1* leads to abnormal development of the DS circuit.

## RESULTS

### Selective deletion of *Isl1* from SACs using *Ptf1a^Cre^*

ISL1 is localized to SACs, ON-BCs, and RGCs in the post-natal retina (Elshatory, Deng, et al., 2007). To better understand the dynamic expression pattern of *Isl1* during retinal development, we used a genetic fate mapping strategy, crossing an *Isl1^Cre^* mouse line to a *Cre*-dependent nuclear *Sun1^LSL-GFP^* reporter **(Fig. S1A**). We labeled SACs with choline acetyltransferase (ChAT) and found that all ChAT^+^ cells were GFP^+^, confirming that all SACs express *Isl1*. Using the AC markers AP-2α and AP-2β (herein called AP2) and another SAC marker (SOX2), we show that no other AC population was labeled in *Isl1^Cre^*;*Sun1^LSL-GFP^* retinas, confirming that SACs are the sole AC subtype that express *Isl1* throughout development (**Fig. S1D, S1E)** (Bassett et al., 2012). Fate mapping using *Isl1^Cre^*;*Sun1^LSL-GFP^* showed that all RGCs (RBPMS^+^) and all BCs (CHX10^+^) express *Isl1* (**Fig. S1B-C**). While ISL1 is localized to ON-BCs but not OFF-BCs in adult mouse retina, the results from fate mapping suggest that OFF-BCs transiently express *Isl1* during early development.

To define the developmental onset of *Isl1* expression in SACs, we crossed *Ptf1a^Cre^* transgenic mice which induce recombination in postmitotic HC and AC precursors with *Cre*-dependent reporter lines (*Ai9*;*R26 ^LSL-TdTomato^* or *Sun1^LSL-GFP^*) and immunostained for ISL1 **(Fig. 1B-C)** (Nakhai et al., 2007). At embryonic day 14.5 (E14.5), when approximately 90% of SACs have been born based on BrdU birthdating, there were very few ISL1^+^/tdT^+^ cells, suggesting that the majority of ISL1^+^ cells are RGCs at this timepoint (Fujitani et al., 2006; Pfaff et al., 1996; Voinescu et al., 2009).

ISL1^+^/tdT^+^ cells were more abundant at E16.5 **(Fig. 1B**), suggesting SACs begin to express *Isl1* approximately two days after they are generated. We then labeled P14 *Ptf1a^Cre^;Sun1^LSL-GFP^* retinas with antibodies against GFP, ISL1, and SOX2, the latter of which marks SACs, astrocytes, and Müller glia (MG) in postnatal retinas (**Fig. 1C**) (Surzenko et al., 2013). All ISL1^+^/SOX2^+^ SACs were GFP^+^, confirming that SACs are the only ISL1^+^ AC subtype and demonstrating that the SAC population can be successfully targeted for recombination by the *Ptf1a^Cre^* line.

Pan-retinal deletion of *Isl1* results in widespread loss of multiple neuronal classes including RGCs, BCs, and SACs (Elshatory, Everhart, et al., 2007). To isolate ISL1’s role specifically in developing SACs without affecting its expression in RGCs and BCs, we generated *Ptf1a^Cre^;Isl1^Flox/+^* control and *Ptf1a^Cre^;Isl1^Flox/Flox^*conditional knockout mice (herein referred to as *Ptf1a^Cre^;Isl1^Ctrl^*and *Ptf1a^Cre^;Isl1^cKO^* mice, respectively). We examined retinas at P14, when SACs have completed their migration, stratified their dendrites in the IPL, and are actively forming synapses (Stacy & Wong, 2003; Wei et al., 2011). In *Ptf1a^Cre^;Isl1^Ctrl^*retinal cross sections, ISL1 labeled SACs, RGCs, and BCs while SOX2 labeled SACs, MG, and astrocytes (**Fig. 1D**). SACs were identified as the only ISL1^+^/SOX2^+^ population, with approximately half of the SACs in the INL (OFF-SACs) and the other half in the GCL (ON-SACs). In P14 *Ptf1a^Cre^;Isl1^cKO^* retinas, ISL1 labeling was maintained in RGCs and BCs while SOX2 labeling was maintained in MG and astrocytes, demonstrating that these populations were unaffected in our conditional deletion strategy. In contrast, very few ISL1^+^/SOX2^+^ cells were observed in the INL and GCL of *Ptf1a^Cre^;Isl1^cKO^* retinas (**Fig. 1D**). These results show that *Isl1* is selectively eliminated from both ON- and OFF-SACs in *Ptf1a^Cre^;Isl1^cKO^* retinas, while its expression in other populations is unaffected.

### Canonical SAC markers are absent in P14 *Ptf1a^Cre^*;*Isl1^cKO^*retinas

To distinguish whether the lack of ISL1^+^/SOX2^+^ cells in *Ptf1a^Cre^;Isl1^cKO^* retinas reflected a loss of SOX2 from putative SACs or a loss of SACs themselves, we examined several SAC-specific and SAC-inclusive markers to determine how they were affected by the deletion of *Isl1* (**Fig. 2**). ChAT and vesicular acetylcholine transporter (VAChT) are direct transcriptional targets of ISL1 in cholinergic neurons in the forebrain and spinal cord and labeled both ON- and OFF-SACs and their dendrites in control retinas (**Fig. 2A, 2B**) (Cho et al., 2014). In contrast, ChAT^+^ and VAChT^+^ labeling was completely absent from P14 *Ptf1a^Cre^;Isl1^cKO^* retinas (**Fig. 2A’-A’’, 2B’-B’’**). We also examined two other SAC molecular markers: Calretinin (CALB2) and Calbindin (CALB1) (Haverkamp & Wässle, 2000). CALB2 is present in multiple RGC and AC subtypes in addition to SACs, labeling dendrites in S2, S3, and S4 of the IPL of control retinas (**Fig. 2C**). *Ptf1a^Cre^;Isl1^cKO^* retinas lacked CALB2^+^ dendrites in S2 and S4 where SACs stratify, but maintained labeling in S3, which corresponds to other ACs and RGCs **(Fig. 2C’-C’’).** CALB1 labeled SAC dendrites in S2/S4 of the IPL and HC dendrites in the OPL of control retinas, but was completely absent in the IPL of *Ptf1a^Cre^;Isl1^cKO^*retinas (**Fig. 2D-D’’**). Taken together, these results show that multiple SAC markers are absent in *Ptf1a^Cre^;Isl1^cKO^*retinas at P14, suggesting that ISL1 plays a critical role in SAC differentiation, survival, and/or maintenance during retinal development.

**Figure 2.**
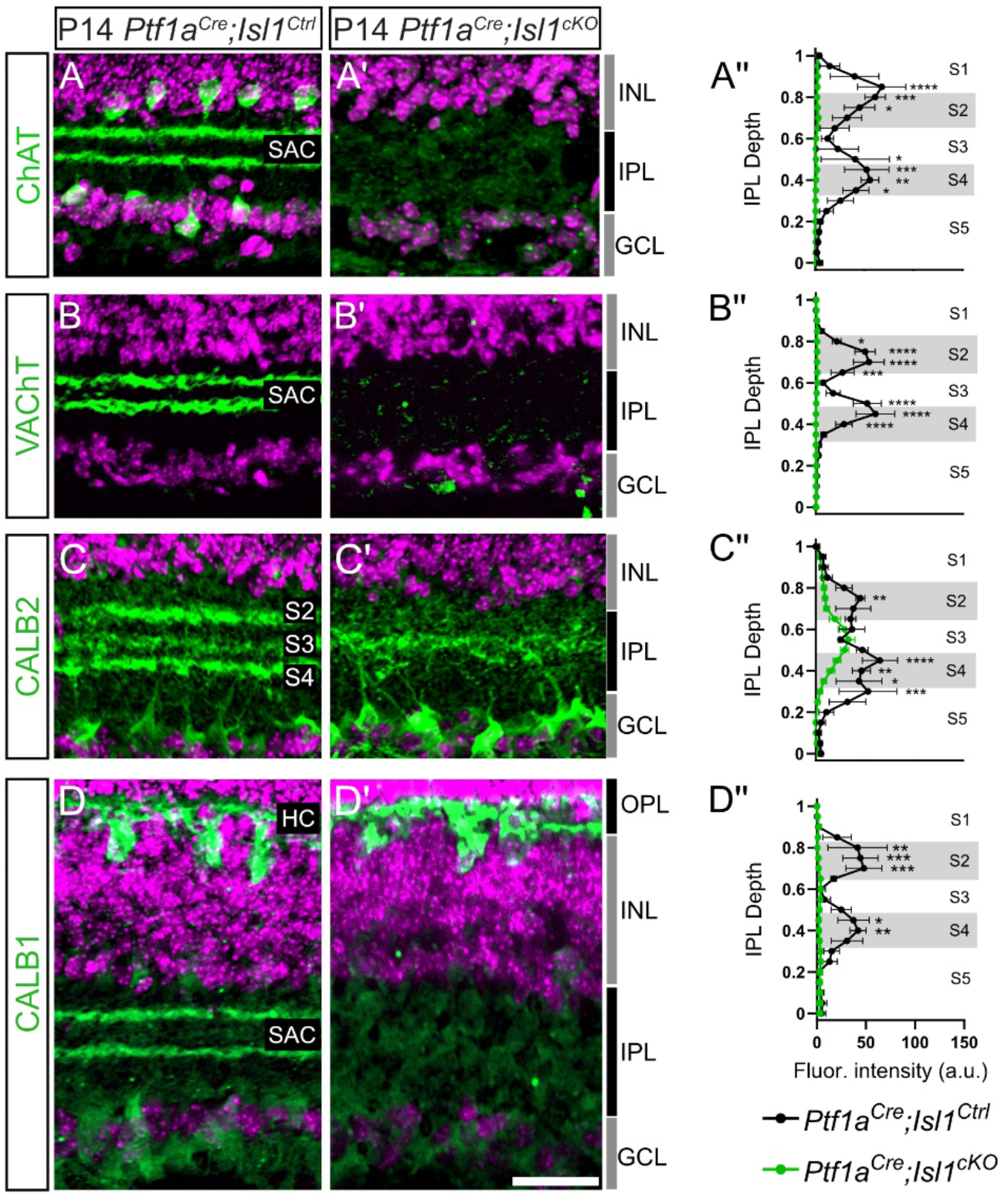
Canonical SAC markers are absent in P14 *Ptf1a^Cre^;Isl1^cKO^* retinas. (**A-D**) Cross sections of P14 *Ptf1a^Cre^;Isl1^Ctrl^* (**A-D**) and *Ptf1a^Cre^;Isl1^cKO^* (**A’-D’**) retinas labeled for ChAT (**A-A’**), VAChT (**B-B’**), CALB2 (**C-C’**), and CALB1 (**D-D’**). Indicated markers are in green, nuclei in magenta. Scale bar = 35µm. (**A’’-D’’**) Fluorescence intensity plots of dendrite stratification. **A-A’’**: n = 2 control; n = 2 mutant. **B-B’’**: n = 3 control; n = 5 mutant. **C-C’’**: n = 3 control; n = 5 mutant. **D-D’’**: n = 3 control; n = 4 mutant. \*\*\*\**p* < 0.0001, \*\*\**p* < 0.001, \*\**p* < 0.01, \**p* < 0.05 by two-way ANOVA with Šidák multiple comparison. Data are represented as mean ± SEM.

### *Isl1^cKO^* SACs are specified but fail to migrate into the GCL

To determine whether ISL1 is required for the initial specification of SACs, we examined *Ptf1a^Cre^;Isl1^cKO^* retinas at earlier developmental timepoints. We first examined P0 retinas when SACs are completing their laminar migration and have separated into distinct layers in the INL and GCL (Ray et al., 2018). Cross-sectioned retinas were immunostained for ISL1 and SOX2 (**Fig. 3A**). In control retinas, ISL1 is only detected in SACs and RGCs as BCs are not born yet, whereas SOX2 is localized to SACs and astrocytes. ISL1^+^/SOX2^+^ SACs were observed in both the INL and GCL at P0 in control retinas. Presumptive SACs that were ISL1^-^/SOX2^+^ were readily identified in *Ptf1a^Cre^;Isl1^cKO^* retinas, however, they were only observed in the INL as opposed to being distributed among the INL and GCL (**Fig. 3A**). Therefore, ISL1 is not required for the initial specification of putative SACs (herein called SOX2^+^ ACs) from AC precursors.

**Figure 3.**
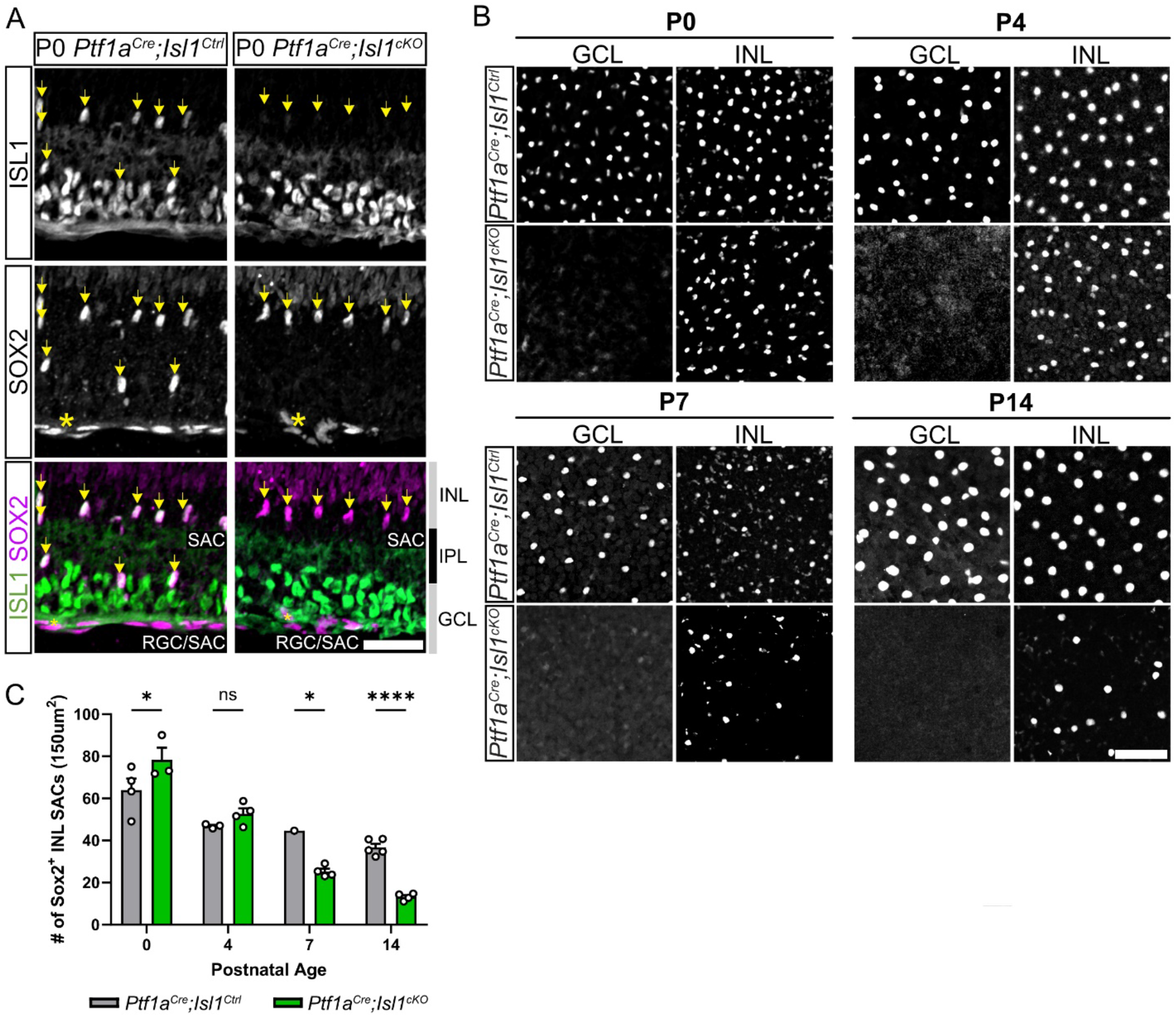
*Isl1^cKO^* SACs are specified but fail to migrate into the GCL. (**A**) P0 *Ptf1a^Cre^;Isl1^Ctrl^* (n = 3) and *Ptf1a^Cre^;Isl1^cKO^* (n = 2) retinas immunostained for ISL1 and SOX2. Yellow arrows indicate SACs (SOX2^+^). Yellow asterisks indicate retinal astrocytes. Scale bar = 35 μm. (**B**) Flatmount retinas of *Ptf1a^Cre^;Isl1^Ctrl^* (top) and *Ptf1a^Cre^;Isl1^cKO^* (bottom) retinas immunolabeled for SOX2 at P0, P4, P7 and P14. Scale bar = 50 μm. (**C**) Counts of INL SOX2^+^ SACs in *Ptf1a^Cre^;Isl1^Ctrl^* and *Ptf1a^Cre^;Isl1^cKO^* at P0 (n = 4, control; n = 3, mutant), P4 (n = 3, control; n = 4, mutant), P7 (n = 1, control; n = 4, mutant), and P14 (n = 5, control, 4, mutant). \*\*\*\**p* < 0.0001, \**p* < 0.05 by two-way ANOVA with Šidák multiple comparison. Data are represented as mean ± SEM.

To understand what happens to SOX2^+^ ACs in *Ptf1a^Cre^;Isl1^cKO^*retinas between P0 when they are present in the INL and P14 where there are very few SOX2^+^ ACs, we analyzed retinal flatmounts at P0, P4, P7, and P14 (**Fig. 3B-C**). In *Ptf1a^Cre^;Isl1^Ctrl^*retinas, SOX2^+^ SACs are present in both the GCL (ON-SACs) and INL (OFF-SACs) at all timepoints. In contrast, SOX2^+^ ACs were only ever observed in the INL at all timepoints in *Ptf1a^Cre^;Isl1^cKO^* retinas (**Fig. 3B-C**). Quantification of SOX2^+^ ACs in the INL across ages revealed that SAC density slightly decreased over time in *Ptf1a^Cre^;Isl1^Ctrl^*retinas, likely reflecting ongoing retinal expansion from P0-P14. The density of SOX2^+^ ACs in *Ptf1a^Cre^;Isl1^cKO^* retinas was significantly increased compared to controls at P0 but steadily decreased from P0 to P14 (**Fig. 3C**). The increased number of SOX2^+^ AC density in the INL of *Ptf1a^Cre^;Isl1^cKO^*retinas at P0 suggests that presumptive SACs are initially specified at approximately the correct numbers, but ON-SACs fail to migrate into the GCL in the absence of *Isl1* (**Fig. 3C**).

### SACs in *Ptf1a^Cre^*;*Isl1^cKO^* retinas undergo BAX-mediated apoptosis

The steady decrease in SOX2^+^ ACs between P0 and P14 in *Ptf1a^Cre^;Isl1^cKO^* retinas could reflect that these cells are either dying or downregulating SOX2 in the absence of *Isl1*. Unlike many other neuronal populations, SACs do not normally undergo developmental apoptosis; deletion of the pro-apoptotic gene *Bax* or overexpression of the pro-survival gene *Bcl2* does not significantly alter SAC numbers (Strettoi & Volpini, 2002; Whitney et al., 2008). To determine whether the gradual loss of SOX2^+^ ACs from P0 to P14 in *Ptf1a^Cre^;Isl1^cKO^* retinas is due to apoptotic cell death, we first examined cleaved-caspase 3 (CC3) at P4 (**Fig. S2**). We did not observe any CC3^+^/SOX2^+^ ACs in either control or mutant retinas in sections or flatmounts at this timepoint. However, because CC3 only provides a snapshot of cells that are actively undergoing apoptosis, we may be unable to detect a gradual loss of SOX2^+^ ACs in *Ptf1a^Cre^;Isl1^cKO^* retinas that occurs over the course of two weeks.

To conclusively test whether *Isl1*-deficient SOX2^+^ ACs are lost due to apoptosis, we crossed a *Bax* knockout line with our *Ptf1a^Cre^;Isl1* line to generate *Bax*-sufficient (*Ptf1a^Cre^;Isl1^Ctrl^;Bax^Ctrl^* and *Ptf1a^Cre^;Isl1^cKO^;Bax^Ctrl^*) and *Bax*-deficient (*Ptf1a^Cre^;Isl1^Ctrl^;Bax^KO^*and *Ptf1a^Cre^;Isl1^cKO^;Bax^KO^*) retinas to either permit or block apoptosis, respectively (**Fig. 4**). P28 retinal cross sections were stained for SOX2 to label SACs and VGLUT3 to label VGlut3-ACs which undergo significant BAX-mediated developmental apoptosis as an internal control (**Fig. 4A, 4E**) (Keeley et al., 2021).

**Figure 4.**
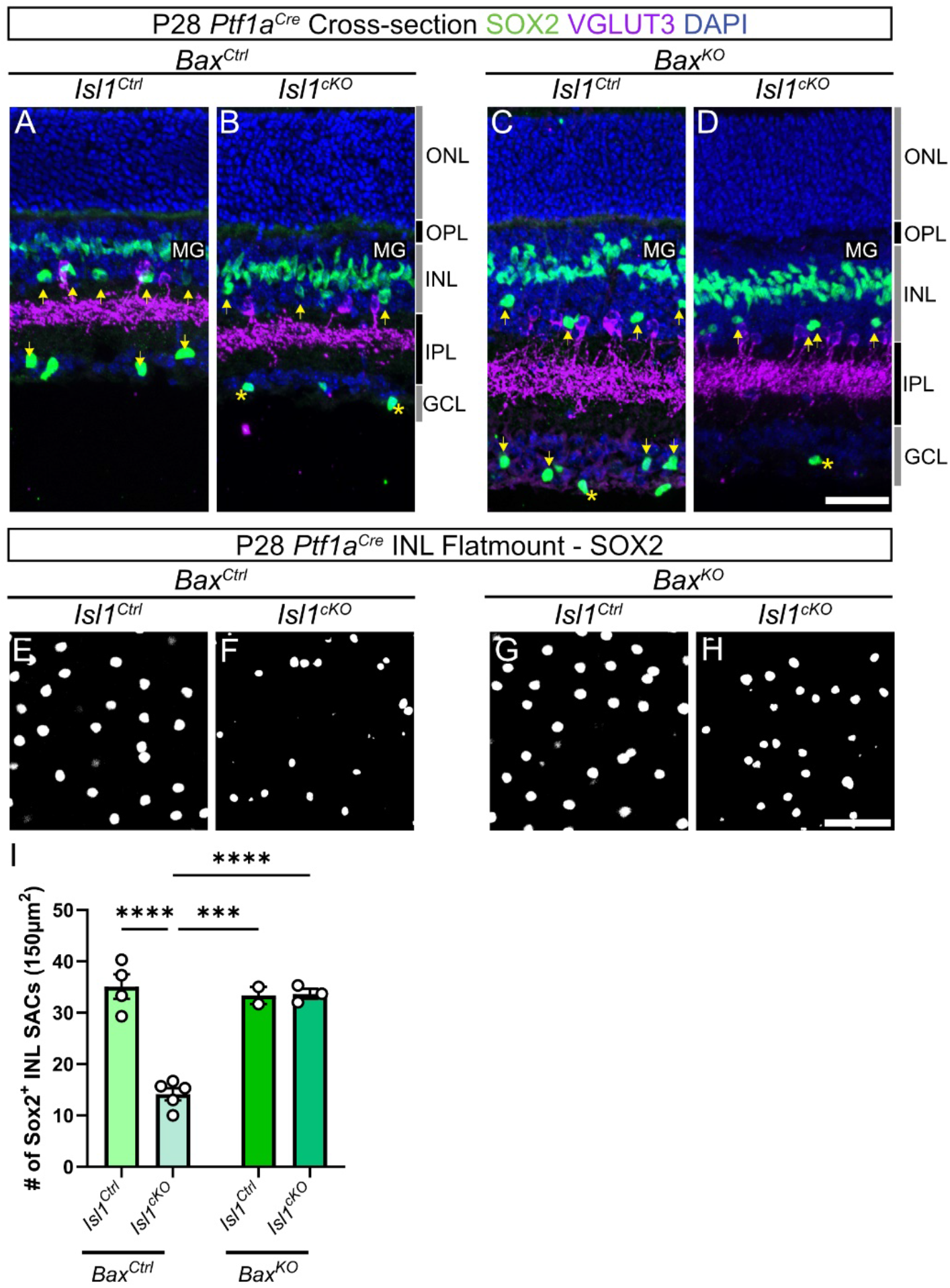
SACs in *Ptf1a^Cre^*;*Isl1^cKO^* retinas undergo BAX-mediated apoptosis. Cross sections of P28 retinas of four genotypes: (**A**) *Ptf1a^Cre^;Isl1^Ctrl^;Bax^Ctrl^*, (**B**) *Ptf1a^Cre^;Isl1^cKO^;Bax^Ctrl^*, (**C**) *Ptf1a^Cre^;Isl1^Ctrl^;Bax^KO^*, and (**D**) *Ptf1a^Cre^;Isl1^cKO^;Bax^KO^* immunostained with SOX2 (green: SACs, Müller glia, astrocytes), VGLUT3 (magenta) and Hoechst (blue). Yellow arrows indicate SACs, yellow asterisks indicate astrocytes. Scale bar = 35 μm. (**E**-**H**) Flatmounts showcasing the INL of the four aforementioned genotypes of P28 retinas. Scale bar = 50 μm. (**I**) Quantification of INL SOX2^+^ SACs in *Ptf1a^Cre^;Isl1^Ctrl^;Bax^Ctrl^* (n = 4), *Ptf1a^Cre^;Isl1^cKO^;Bax^Ctrl^* (n = 5), *Ptf1a^Cre^;Isl1^Ctrl^; Bax^KO^* (n = 2), *Ptf1a^Cre^;Isl1^cKO^;Bax^KO^*(n = 3). \*\*\*\**p* < 0.0001, \*\*\**p* < 0.001 by two-way ANOVA with Tukey multiple comparison. Data are represented as mean ± SEM.

Overall retinal thickness increased in *Ptf1a^Cre^;Isl1^Ctrl^;Bax^KO^*and *Ptf1a^Cre^;Isl1^cKO^;Bax^KO^* retinas due to the inability of many retinal neurons to undergo normal levels of BAX-mediated developmental apoptosis. SACs in *Ptf1a^Cre^;Isl1^Ctrl^*retinas were unaffected by the loss of *Bax*, whereas the VGlut3-ACs dendrite stratification in the IPL was thicker due to the increased number of VGlut3-ACs (**Fig. 4C, 4G**). *Ptf1a^Cre^;Isl1^cKO^;Bax^Ctrl^* retinas recapitulated the phenotype of *Ptf1a^Cre^;Isl1^cKO^* retinas, with only a few SOX2^+^ ACs remaining exclusively in the INL **(Fig. 4B, 4F**). In the double mutant retinas (*Ptf1a^Cre^;Isl1^cKO^;Bax^KO^*), SOX2^+^ ACs were present but were exclusively located in the INL (**Fig. 4D, H**). Quantification of SOX2^+^ ACs in the INL across all four conditions at P28 revealed that the density in the INL of *Ptf1a^Cre^;Isl1^cKO^;Bax^KO^*is restored to similar levels seen in control conditions (**Fig. 4I**). Therefore, a significant proportion of SOX2^+^ ACs undergo BAX-mediated apoptosis in the absence of *Isl1*.

### ISL1 is required for SAC terminal differentiation

The absence of SAC markers at P14 (**Fig. 2**) likely reflects the near complete loss of SACs themselves due to BAX-mediated apoptosis (**Fig. 4**). We therefore examined pan-AC and SAC-specific markers at P4 when the number of INL SOX2^+^ ACs is roughly equivalent across genotypes (**Fig. 3**). To broaden our selection of SAC-specific markers beyond the cholinergic battery, we referenced single-cell RNA-sequencing datasets from SACs at E16 and P0 and included *FRMD7* and *Gng7* as additional SAC-specific genes (Peng et al., 2020). Pan-AC markers (AP2, STX1A, PAX6, GAD67) were unaffected by the loss of *Isl1*, with all SOX2^+^ ACs remaining double positive in control and *Ptf1a^Cre^;Isl1^cKO^* retinas (**Fig. 5A-D, I**). Conversely, the SAC-specific markers CALB1, CALB2, VAChT, ChAT, *FRMD7*, and *Gng7* were completely absent from SOX2^+^ ACs in *Ptf1a^Cre^;Isl1^cKO^* retinas (**Fig. 5E-I, Fig. S3**). Three other SAC-specific markers (INA, MEGF10, and *Fezf1*) were minimally perturbed by the loss of *Isl1* in mutant SOX2^+^ ACs, demonstrating that some SAC-specific genes do not require ISL1 (**Fig. S4**). Overall, the loss of SAC-specific markers but not pan-AC markers are consistent with ISL1’s role in directing the terminal differentiation of SACs from a generic amacrine identity to a SAC-specific one.

**Figure 5.**
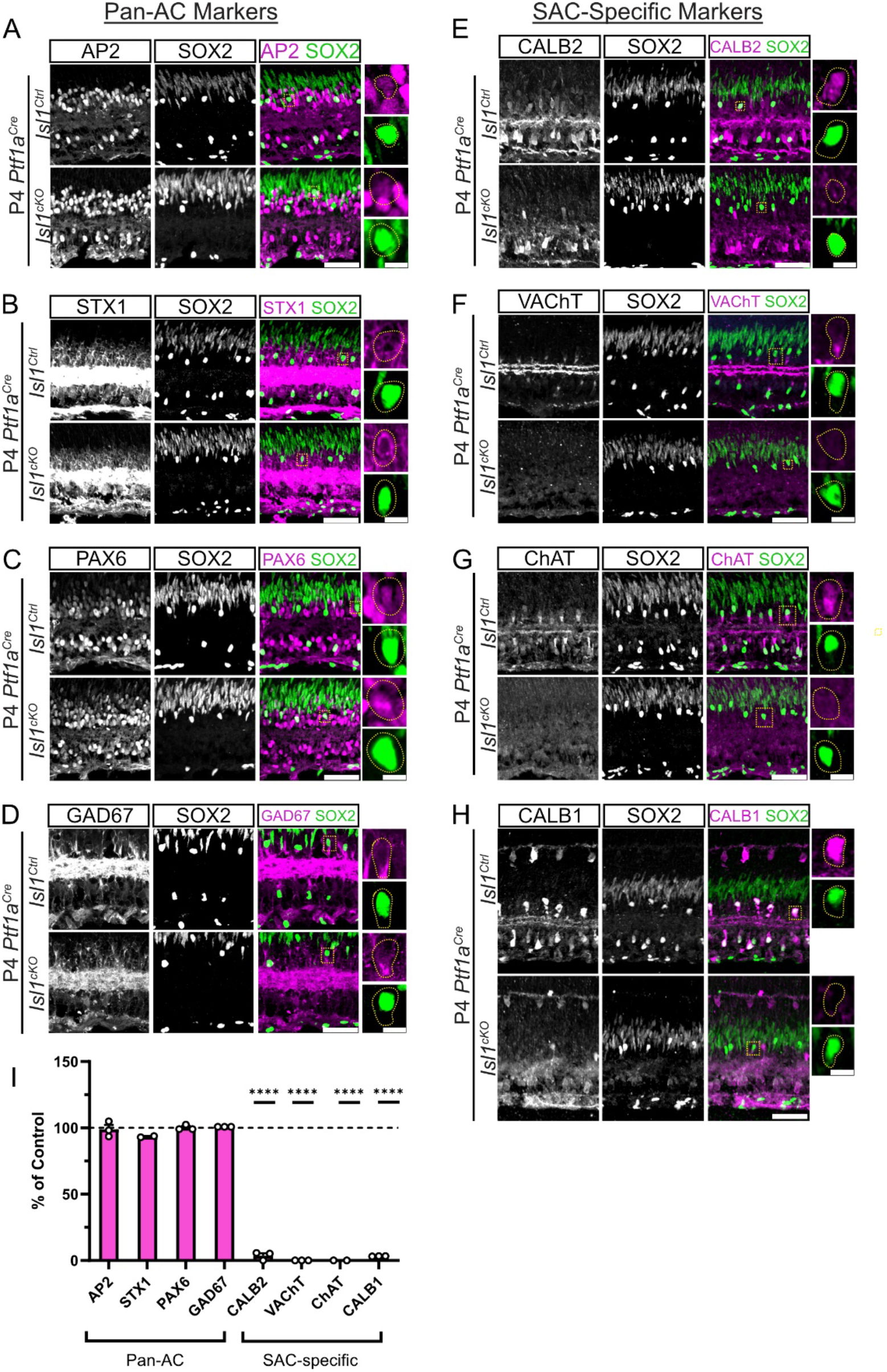
ISL1 is required for SAC terminal differentiation. (**A-D**) Cross section of P4 *Ptf1a^Cre^;Isl1^Ctrl^* (top) and P4 *Ptf1a^Cre^;Isl1^cKO^* (bottom) retinas labeled with pan-AC markers (magenta: **A**: AP2, **B**: STX1A, **C**: PAX6, **D**: GAD67) and SAC-marker SOX2 (green) with corresponding insets. (**E-H**) Cross section of P4 *Ptf1a^Cre^;Isl1^Ctrl^* (top) and P4 *Ptf1a^Cre^;Isl1^cKO^* (bottom) retinas labeled with SAC-specific markers (magenta: **E**: CALB2, **F**: VAChT, **G**: ChAT, **H**: CALB1) and SAC-marker SOX2 (green) with corresponding insets. Scale bar = 50μm. Inset scale bar = 7.5μm. (**I**) Quantification of SACs (SOX2^+^) positive for pan-amacrine markers (AP2, n = 3; STX1A, n = 2; PAX6, n = 3; GAD67, n = 3) and SAC-specific markers (CALB2, n = 3; VAChT, n = 3; ChAT, n = 2; CALB1, n = 3). Quantification of SOX2^+^ cells that were marker^+^ in mutants was normalized to littermate controls (n = 6). \*\*\*\**p* < 0.0001, by two-way ANOVA with Šidák multiple comparison. Data are represented as mean ± SEM.

### ISL1 Is required to maintain SAC identity

In addition to directing the acquisition of subtype-specific effector genes as neuronal subtypes differentiate, some transcription factors are also required for the continuous maintenance of neuronal identity (Deneris & Hobert, 2014). Therefore, we generated *ChAT^Cre^;Isl1^cKO^;R26^LSL-tdTomato^* mice to delete *Isl1* from SACs at a later stage of differentiation while simultaneously genetically labeling them with tdTomato (**Fig. 6**). *ChAT^Cre^* has been commonly used to assess SAC physiology (Kozlowski et al., 2024; Pei et al., 2015; Ray et al., 2018; Xu et al., 2016). *ChAT^Cre^* recombination occurs in SACs during the first post-natal week, several days after SAC generation and migration to the inner retina (Xu et al., 2016). *ChAT^Cre^;Isl1^cKO^*mice were small and died within the first two postnatal weeks due to apparent respiratory failure, reflecting defects in cholinergic neurons outside the retina (**Fig. S5A**). This limitation prompted us to collect retinas at P12. We first analyzed SAC survival in flatmount by counting tdT^+^ cells. In contrast to *Ptf1a^Cre^;Isl1^cKO^* retinas, survival of SACs was unaffected in *ChAT^Cre^;Isl1^cKO^* retinas at P12 (**Fig. 56B-C**). Cross sections were assessed for pan-AC markers (AP2, PAX6, GAD67, and STX1A) and SAC-specific markers (CALB1, CALB2, SOX2, ChAT and VAChT). All pan-AC markers were unaffected in *ChAT^Cre^;Isl1^cKO^* retinas (**Fig. 6A-D, 6J**), similar to what we observed with the earlier deletion of *Isl1* in *Ptf1a^Cre^;Isl1^cKO^* retinas (**Fig. 5**). Of the SAC-specific markers, SOX2 was unaffected, CALB1 and CALB2 were completely absent in P12 *ChAT^Cre^;Isl1^cKO^* retinas, while ChAT and VAChT were present but significantly reduced (**Fig. 6E-I, 6K-L**). We also noted that SAC dendrite stratification was abnormal in *ChAT^Cre^;Isl1^cKO^* retinas; tdTomato^+^ dendrites were highly disorganized and frequently strayed out of the normally tight bands in S2 and S4 (**Fig 6E-I, 6K**). This data shows that in addition to its role in directing SAC terminal differentiation, ISL1 is required to maintain SAC identity and proper dendrite stratification.

**Figure 6.**
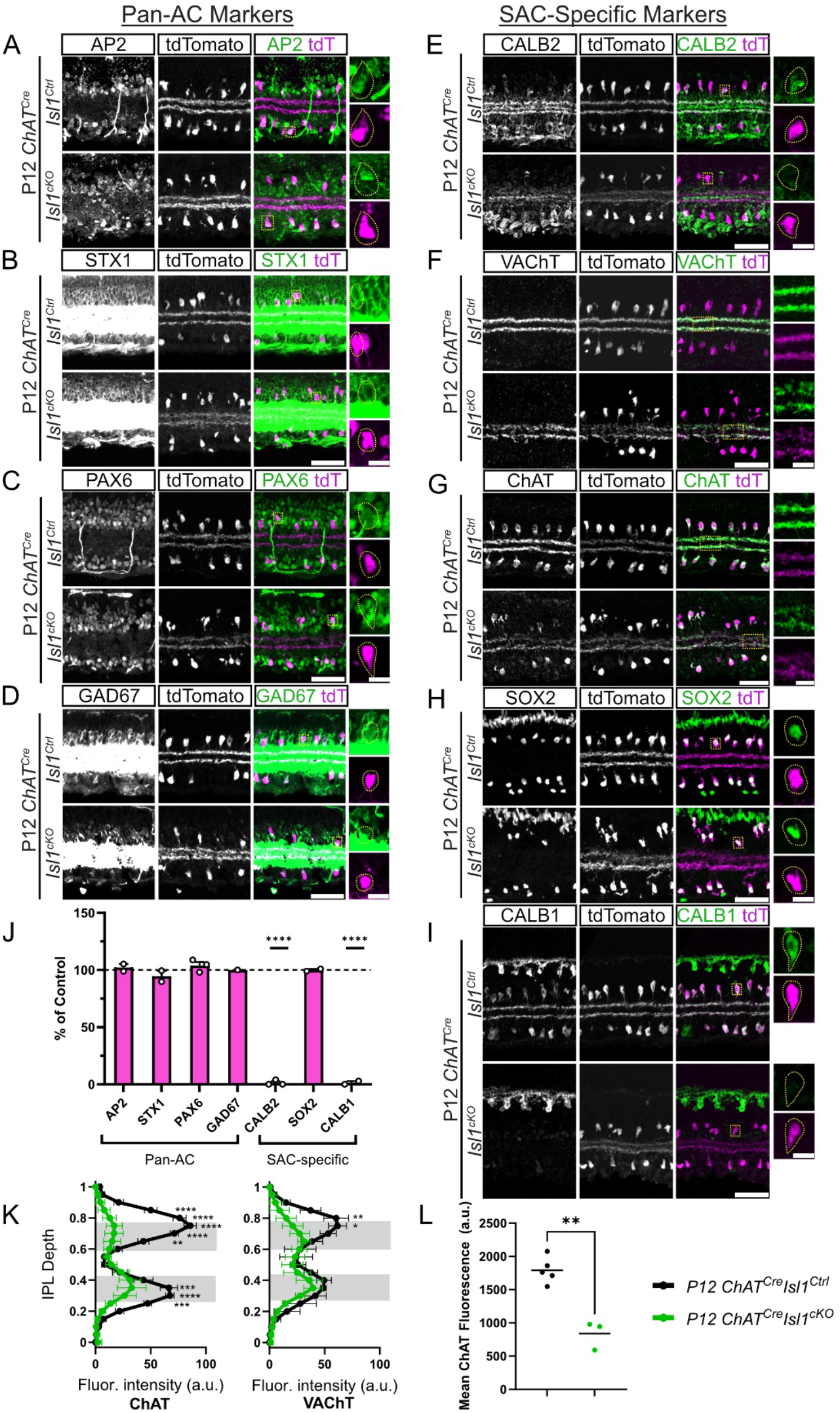
ISL1 is required to maintain SAC identity. (**A-I**) Cross section of P12 *ChAT^Cre^;Isl1^Ctrl^* (top) and P12 *ChAT^Cre^;Isl1^cKO^* (bottom) retinas labeled with pan-AC markers (magenta) (**A**) AP2, (**B**) STX1A, (**C**) PAX6 and (**D**) GAD67 or SAC-specific markers (**E**) CALB2, (**F**) VAChT, (**G**) ChAT, (**H**) SOX2 and (**I**) CALB1 with corresponding insets. SACs are labeled with tdT (magenta). Scale bar = 50μm. Inset scale bar = 7.5μm. (**J**) Quantification of tdT^+^ SACs with corresponding markers from (**A-I**) Quantification of mutants is normalized to littermate controls (n = 5). (AP2, n = 2; STX1A, n = 2; PAX6, n = 3; GAD67, n = 1; CALB2, n = 3; SOX2, n = 2; CALB1, n = 2). (**K)** IPL fluorescence analysis of ChAT (control, n = 5; mutant, n = 3) and VAChT (control, n = 5; mutant, n = 3). (**L**) ChAT cell body fluorescence analysis between control (n = 5) and mutant (n = 3). \*\*\*\**p* < 0.0001, \*\*\**p* < 0.001, \*\**p* < 0.01, \**p* < 0.05. (**J & K**) calculated by two-way ANOVA with Šidák multiple comparison. (**L**) calculated by unpaired Welch’s t-test. All data are represented as mean ± SEM.

### Loss of *Isl1* from SACs selectively affects the development of the direction selective circuit

Pan-retinal deletion of *Isl1* using *Six3^Cre^* results in the loss of multiple retinal neuron types including SACs, other ACs, RGCs, and BCs (Elshatory, Everhart, et al., 2007). Pan-retinal deletion of *Isl1* also results in an increase in HCs, which do not express *Isl1* in the mouse retina based on fate mapping results (**Fig. S1**) (Elshatory, Everhart, et al., 2007). We found that HC numbers were unaffected in *Ptf1a^Cre^;Isl1^cKO^*retinas (**Fig. S6A-B**), excluding the possibility that SOX2^+^ ACs are fate switching to HCs in the absence of *Isl1*. We next examined whether the disruptions in SAC development affected other AC populations. Staining for dopaminergic (TH^+^) and VGlut3 (VGLUT3^+^) AC populations showed no difference between the dendritic stratification pattern of either AC subtypes in control or *Ptf1a^Cre^;Isl1^cKO^* retinas (**Fig. S6C-F**).

SACs are part of the DS circuit, receiving synaptic inputs from multiple OFF- and ON-BC subtypes and providing synaptic output to the four cardinal ON-OFF DS RGC subtypes (ooDSGCs) and the three ON DS RGC subtypes (oDSGCs) (**Fig. 7A**) (Briggman et al., 2011; H. Ding et al., 2016; Duan et al., 2014; Fried et al., 2002; Trenholm et al., 2011). Previous work has shown that SACs play a key role in the assembly of the DS circuit during development. Mis-stratifying SAC dendrites can recruit their synaptic partners to inappropriate layers in the IPL, and genetic ablation of SACs during early development causes abnormal stratification of ooDSGCs (Duan et al., 2018; Prigge et al., 2023; Ray et al., 2018). To examine how inappropriate migration and subsequent loss of SACs in *Ptf1a^Cre^;Isl1^cKO^* mice affects the development of the DS circuit, we used a transgenic *Hb9-GFP* reporter line that labels ventral-preferring ooDSGCs. GFP^+^ ooDSGC cell counts showed no difference between control and in *Ptf1a^Cre^;Isl1^cKO^* mutants, indicating that ooDSGC survival is unaffected by the loss of SACs at postnatal ages (**Fig. 7D**). In P14 *Ptf1a^Cre^;Isl1^Ctrl^;HB9^GFP^*retinas, GFP^+^ ooDSGC dendrites were bistratified in S2 and S4. In contrast, GFP^+^ ooDSGC dendrites in *Ptf1a^Cre^;Isl1^cKO^* retinas only stratified in S2 with no detectable S4 stratification (**Fig. 7B-C**). This suggests that the SOX2^+^ ACs present in the INL of *Ptf1a^Cre^;Isl1^cKO^* mutants likely stratify their dendrites in S2 at early timepoints prior to undergoing apoptosis. Synaptotagmin-2 (SYT2) labeling of Type 2 and Type 6 BC axon terminals did not show any difference between control and *Ptf1a^Cre^;Isl1^cKO^* retinas, indicating that stratification of pre-synaptic BCs is unaffected by the loss of *Isl1* from SACs (**Fig. 7E-F**).

**Figure 7.**
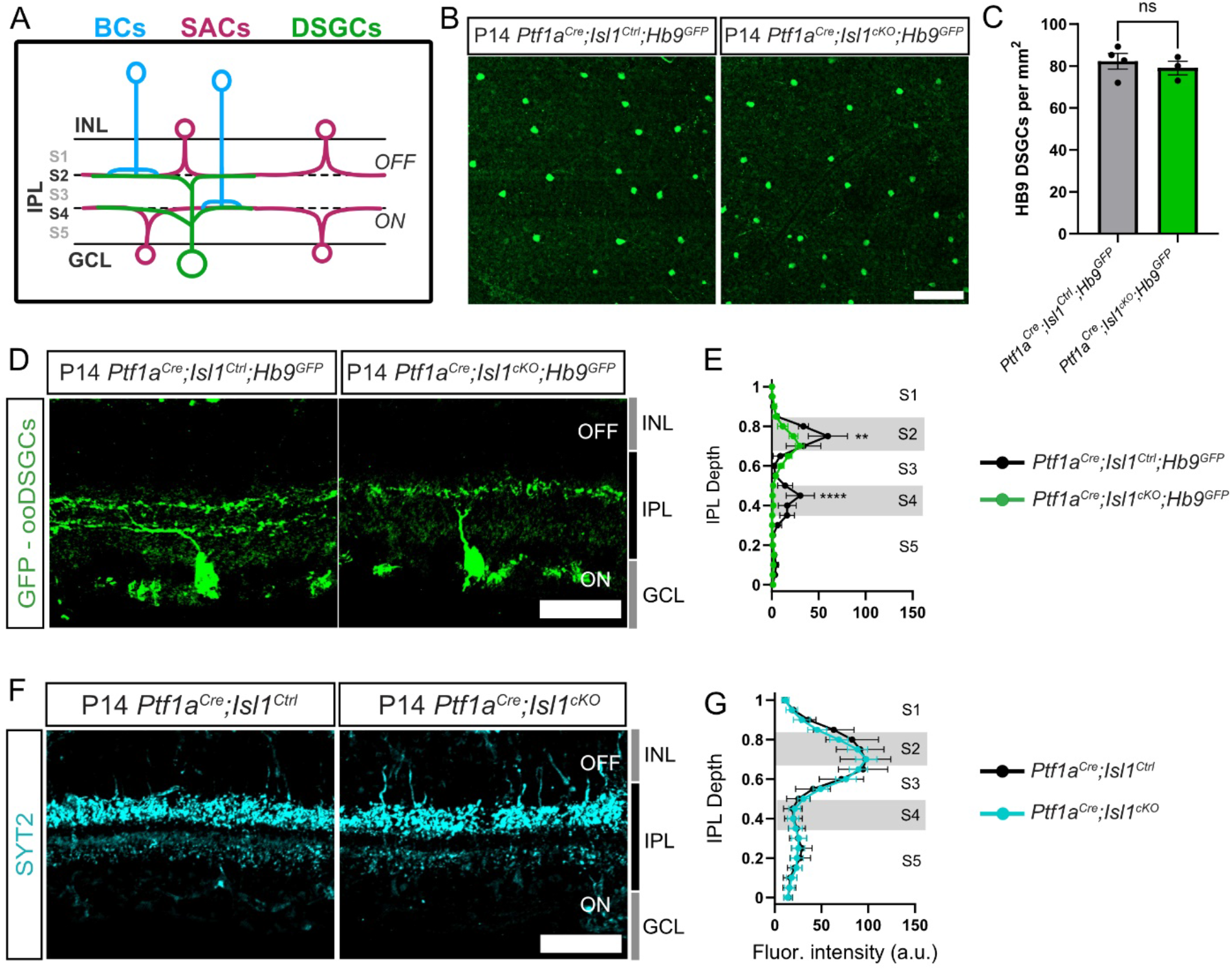
Loss of *Isl1* from SACs disrupts the development of the direction selective circuit. (**A**) Schematic of the DS circuit containing BCs, SACs, and ooDSGCs costratifying in S2 and S4 of the IPL. (**B**) Flatmount of P28 *Ptf1a^Cre^Isl1^Ctrl^Hb9^GFP^* (left) and *Ptf1a^Cre^Isl1^cKO^Hb9^GFP^*(right) retinas immunolabeled for GFP to label Hb9-ooDSGCs. Scale bar = 100µm. (**C**) Quantification of Hb9^+^ ooDSGC cell bodies from (**B**) (control, n = 4; mutant, n = 3). (**D**) Cross sections of P14 *Ptf1a^Cre^Isl1^Ctrl^Hb9^GFP^* (left) and *Ptf1a^Cre^Isl1^cKO^Hb9^GFP^*(right) retinas immunolabeled with GFP to label Hb9-ooDSGC dendrites. Scale bar = 50µm. **(E**) Mean intensity (±SEM) of GFP^+^ ooDSGC dendritic bands in P14 *Ptf1a^Cre^Isl1^Ctrl^Hb9^GFP^* (n = 3) and *Ptf1a^Cre^Isl1^cKO^Hb9^GFP^* (n = 4). Scale bar = 50µm. (**F**) Cross sections of P14 *Ptf1a^Cre^;Isl1^Ctrl^*(left) and *Ptf1a^Cre^;Isl1^cKO^* retinas immunolabeled with BC synaptic marker SYT2. Scale bar = 50µm. (**G**) Mean intensity (±SEM) of (**F**) in P14 *Ptf1a^Cre^;Isl1^Ctrl^*(n = 3) and *Ptf1a^Cre^;Isl1^cKO^* (n = 5).

## DISCUSSION

The recent development of single-cell transcriptomics has resulted in a dramatic increase in the description of molecularly distinct neuronal subtypes, raising the question of how these subtypes emerge during development. To understand the transcriptional logic that drives the differentiation and maintenance of individual neuronal subtypes, careful and precise genetic manipulation of molecular regulators is required. Here we developed an approach that allowed us to conditionally delete *Isl1* from HC and AC precursors to examine ISL1’s cell-autonomous role in SAC development without secondary effects elsewhere in the retina. We show that ISL1 is dispensable for the initial specification of SACs. However, ISL1 is critical for the terminal differentiation and maintenance of SACs from a generic AC identity. *Isl1*-defiicent SACs fail to express canonical SAC markers and show aberrant migration and dendrite stratification patterns, ultimately leading to the abnormal development of the DS circuit.

### ISL1’s cell-autonomous role in SACs

Previous results found that SACs are largely absent following pan-retinal deletion of *Isl1* (Elshatory, et al., 2007). However, it was unclear whether this reflected a cell-autonomous defect in SAC specification, terminal differentiation, or maintenance due to two main caveats. First, pan-retinal deletion of *Isl1* results in the loss of many other inner retinal neuron types, including pre- and post-synaptic partners of SACs. Affected populations also included OFF-BCs, Müller glia, and HCs which do not express *Isl1* in the adult retina, suggesting that some of these effects may be indirect. Second, the markers used to identify SACs in this prior study (ChAT, CALB2) are no longer expressed in SACs in the absence of ISL1 (**Figs. 2, 5**) and are likely to be direct transcriptional targets of ISL1. Our approach of selectively deleting *Isl1* from AC precursors allowed us to circumvent any non-cell autonomous effects by preserving expression of *Isl1* in RGCs and ON-BCs. Using SOX2 as a marker, we found that SACs are present in *Ptf1a^Cre^;Isl1^cKO^* at early postnatal timepoints, showing that SACs are initially specified in the absence of *Isl1* (**Fig. 3**). These SOX2^+^ SACs continue to express pan-AC markers but lose expression of many canonical SAC-specific genes (**Fig. 5**). Subsequent deletion of *Isl1* from post-migratory SACs using *ChAT^Cre^* also resulted in a loss of many SAC-specific markers (**Fig. 6**). Therefore, *Isl1* appears to be required for both the terminal differentiation and maintenance of SACs from a more generic AC identity, but not for their initial specification.

Currently, we have not identified the full complement of dysregulated genes downstream of ISL1 in SACs. However, previous work has shown that the entire cholinergic gene battery is a direct transcriptional target of ISL1 in other types of cholinergic neurons (Cho et al., 2014). In spinal cord motor neurons and forebrain cholinergic neurons, ISL1 forms hexameric complexes with NLI and LHX3 or LHX8, respectively. These complexes directly bind hexamer response elements in *ChAT*, *VAChT*, *Slc5a7*, and *Acly*, which are all required for the synthesis and packaging of acetylcholine, establishing the cholinergic gene battery in two developmentally unrelated neuronal populations. While the selective expression of *ISL1* and *NLI* in cholinergic amacrine cells is evolutionarily conserved from mice to primates, SACs do not express appreciable levels of any *LHX* family member (Kay et al., 2011; Peng et al., 2020; Tommasini et al., 2026). This suggests while ISL1 is likely to play a conserved role in the terminal differentiation of vertebrate SACs, it may use distinct co-factors compared to other cholinergic neuron populations.

### Parallel transcriptional programs regulate SAC migration

While loss of *Isl1* prevents proper terminal differentiation of SACs, a subset of SAC-specific markers are retained in the absence of *Isl1* including SOX2, *Fezf1*, INA, and MEGF10 suggesting that they are controlled by parallel pathways independent of ISL1 (**Fig. S4**). Of particular interest are SOX2 and *Fezf1*, which both play a role in SAC migration without affecting *Isl1* expression. Conditional deletion of *Sox2* from SACs using *ChAT^Cre^* results in a modest reduction in the number of SACs that migrate into the GCL and abnormal dendrite stratification in a subset of ON-SACs (Whitney et al., 2014).

It is likely that these relatively mild phenotypes are due to the timing of *Sox2* deletion with *ChAT^Cre^*, and that earlier deletion of *Sox2* with *Ptf1a^Cre^* would result in more severe migration defects. *Fezf1* is selectively expressed in ON-SACs, where it represses OFF-SAC identity and controls the migration of ON-SACs into the GCL (Peng et al., 2020). *Fezf1^-/-^* retinas show a fully penetrant SAC migration phenotype, where all SACs remain in the INL, similar to what we observed in *Ptf1a^Cre^;Isl1^cKO^*retinas. The similarities between *Isl1*, *Sox2*, and *Fezf1* mutant SACs suggest that these distinct transcription factors control parallel pathways that regulate ON-SAC migration into the GCL. While SOX2 and FEZF1 both appear to regulate SAC migration, they do not appear to regulate overall terminal differentiation of SACs, as both *Sox2* and *Fezf1* mutant SACs retain their cholinergic identity and many other SAC-specific genes (Peng et al., 2020; Whitney et al., 2014)

### SACs as the scaffold for the DS circuit

Previous work has shown that SACs play a key role in the formation of the DS circuit, acting as a physical scaffold for their synaptic partners. This is thought to occur in part because SACs stratify their dendrites in the IPL before the rest of their DS circuit counterparts. Genetic manipulations that specifically affect SAC dendritic lamination cause mis-stratification of ooDSGCs and defective DS circuit function (Duan et al., 2018; Peng et al., 2017; Prigge et al., 2023; Ray et al., 2018; Stacy & Wong, 2003). Our results add to this body of work: we find that in *Ptf1a^Cre^;Isl1^cKO^;Hb9^GFP^* retinas, the GFP^+^ ooDSGCs monostratify in S2 instead of stratifying in both S2 and S4. While we are unable to label SAC dendrites with ChAT or VAChT in *Ptf1a^Cre^;Isl1^cKO^*retinas, staining with INA clearly shows mutant SACs stratifying in S2 of the INL (**Fig. S4A**). These results suggest that the SACs that are present in the INL of *Ptf1a^Cre^;Isl1^cKO^* mutants at early postnatal ages serve as a scaffold for the ooDSGC dendrites. However, we are unable to assess this directly due to the lack of markers that label SAC morphology in these mutants.

### Transcriptional regulation of cholinergic differentiation across species

While the expression of *ISL1* in distinct subtypes of cholinergic neurons is highly conserved in mammals, the *ISL1* homologues *tailup* and *lim-7* are not expressed in cholinergic neurons in *Drosophila* or *C. elegans*, respectively (Certel & Thor, 2004; Wolfram et al., 2014). Instead, other transcription factors appear to regulate cholinergic identity in these invertebrate species. In *Drosophila*, Acj6 directly binds to the *chat* locus in a subset of cholinergic neurons and ChAT levels are decreased in *acj6* mutants (Estacio-Gómez et al., 2020; Lee & Salvaterra, 2002). *Apterous* is expressed in cholinergic neurons in the *Drosophila* optic lobe, where it is required to maintain the expression of *chat* in most but not all of these neurons; whether *apterous* acts directly to regulate the cholinergic gene battery is unknown. In the *C. elegans* ventral nerve cord, *unc-3* cooperates with an array of co-factors to directly regulate expression of the cholinergic gene battery in ventral nerve cord motor neurons and the SAB head motor neuron. Other transcription factors (*unc-42*, *lim-4*, *lin-11*, *unc-30*, *unc-86*, and *ceh-14*) regulate cholinergic identity in other neuronal subtypes (Pereira et al., 2015; Zhang et al., 2014). Similar to our findings in *Isl1*-deficient SACs, loss of these transcription factors in both *Drosophila* and *C. elegans* does not affect the expression of pan-neuronal genes (Kratsios et al., 2015; Pereira et al., 2015; B. Prasad et al., 2008; B. C. Prasad et al., 1998; Zhang et al., 2014). Interestingly, *Isl1* and many of the transcription factors that regulate cholinergic identity in *Drosophila* (*apterous*) and *C. elegans* (*lim-4*, *lin-11*, *ceh-14*) are members of the LIM homeodomain family, suggesting evolutionary conservation. One reason that *Drosophila* and *C. elegans* may have multiple, non-redundant transcription factors for directing cholinergic identity is that over half of their neurons release acetylcholine, making it their most widely used neurotransmitter (Croset et al., 2018; Pereira et al., 2015).

### Neuronal subtype specific terminal differentiation in the mammalian nervous system

Our data show that the requirement for ISL1 changes over the course of SAC development. Embryonic deletion of *Isl1* from SACs with *Ptf1a^Cre^* results in their failure to undergo terminal differentiation and subsequent apoptotic cell death during the first two postnatal weeks. In contrast, *Isl1* deletion from post-migratory SACs with *ChAT^Cre^* results in a loss of SAC markers without affecting their survival, at least until the latest timepoint we were able to examine (P12). These temporal distinctions of ISL’s role in regulating terminal differentiation, maturation, and survival in SACs parallel a recent study examining the transcription factor Gfi1 in two RGC subtypes: downward preferring-ON-DSGCs and F-mini-ONs (Chaudhari et al., 2026).When *Gfi1* is constitutively deleted, these RGC subtypes fail to differentiate normally, and a large proportion are lost between P1 and P6. Conditional deletion with *CreERT2* lines to provide temporal control reveal an ongoing requirement for *Gfi1* in the survival of F-mini-ONs, with deletion at P0 or P40 resulting in the loss of these cells. In contrast, postnatal deletion of *Gfi1* affects the morphology and function of downward preferring-ON-DSGCs, but is not required for their survival.

There are also temporally distinct roles for transcription factors involved in the terminal differentiation and maintenance of dopaminergic and serotonergic neurons in the brain. Transcription factors required for dopaminergic neuron terminal differentiation include *Etv1*, *Pbx1*, *Pitx3,* and *Nurr1* (Flames & Hobert, 2009; Kadkhodaei et al., 2009; K.-S. Kim et al., 2003; Remesal et al., 2020), while *Pet1* and *Lmx1b* are required in serotonergic neurons (Y.-Q. Ding et al., 2003; Hendricks et al., 2003; Krueger & Deneris, 2008; C. Liu et al., 2010; Zhao et al., 2006). In each of these cases, the deletion of the specific transcription factor results in a loss of dopaminergic or serotonergic identity while pan-neuronal features are maintained. However, the loss of these transcription factors has differing effects on the survival of their target neuron populations depending on the context. For example, while neurons fail to adopt a serotonergic identity in *Pet1* mutants, these partially differentiated neurons persist into adulthood (Krueger & Deneris, 2008). When *Pbx1* is conditionally deleted from dopaminergic neurons in the olfactory bulb, these neurons fail to adopt a dopaminergic identity but are able to survive long-term (Remesal et al., 2020). In contrast, when *Lmx1b* is conditionally deleted early from serotonergic populations, the neurons are initially specified at the correct number but are gone by adulthood presumably due to cell death (Zhao et al., 2006). Deletion of *Pet1* or *Lmx1b* from adult serotonergic neurons or *Nurr1* from dopaminergic neurons results in the loss of neurotransmitter identity and degeneration of synaptic terminals and axons, but the neurons survive (Kadkhodaei et al., 2013; Kitt et al., 2022). Overall, whether the transcription factors required for the initial terminal differentiation of a neuronal subtype are required for their long-term survival and/or maintenance of neuronal identity appears to be case-dependent.

Taken together, our results identify a critical role for ISL1 in the acquisition and maintenance of SAC-specific features from a generic AC identity. Future work will be needed to define a more comprehensive view of ISL1 effector genes in SACs, particularly those involved in regulating their migration and dendrite stratification. ISL1 likely plays a similar role in maintaining the identity of other cholinergic populations in the brain and spinal cord, although more precise genetic manipulations will be needed to test this directly. Ultimately, these findings add to the growing body of work showing how transcription factors can control the development and maintenance of neuronal subtypes in the mammalian nervous system.

## Methods and Materials

### Animal husbandry

All animals were housed and cared for by the Department of Comparative Medicine (DCM) at Oregon Health & Science University (OHSU), an AAALAC-accredited institution. All animal usage, care, and procedures were approved by the OHSU Institutional Animal Care and Use Committee (Protocol # IS00000539), adhered to the NIH *Guide for the care and use of laboratory animals* and provided with 24-hour veterinary care. Mouse facilities were regulated for temperature, humidity and maintained on a 12-hour light-dark cycle and provided food and water *ad libitum.* All lines were maintained on a *C57BL/6J* background and mice of both sexes were used for experiments. Mice were used between aged E14.5 and P28 as indicated in the text or figure legend. Mice younger than P10 were euthanized by decapitation whereas mice older than P10 were euthanized by CO_2_ inhalation followed by cervical dislocation. For all experiments and timepoints, a minimum of three mice per genotype from at least two independent litters were analyzed. Mutant mice were compared to littermate controls for all experiments and timepoints with the exception of Figures 4 and 7 due to low probability of obtaining all relevant genotypes in a single litter.

### Mouse strain and genotyping

The day of birth was designated as postnatal day 0 (P0). Specific ages of mice used for each experiment is indicated in both the figure and figure legend. Male *Ptf1a^Cre^* mice were bred to female homozygous *Isl1^Flox^* mice to generate *Ptf1a^Cre^;Isl1^Flox/+^*male breeders. Experimental mice were obtained by pairing *Ptf1a^Cre^;Isl1^Flox/+^*mice with homozygous *Isl1^Flox^* female mice with and without reporters (tdTomato or Sun1^GFP^).

*ChAT^Cre^;Isl1^Flox/+^* males were paired with homozygous *Isl1^Flox^;R26^tdTomato^* female mice to obtain experimental mice. Toe or tail samples were used to extract genomic DNA using the HotSHOT method to genotype animals. The following primers were used to genotype *Isl1^Flox^*mice: TCTCTCATTGGGGAAGCAAC (flox forward), and CCAGTGAAGGCCTTCCAGT (flox reverse). *Sun1^LSL-GFP^* mice were genotyped using the following primers: AAGGGAGCTGCAGTGGAGTA (WT forward), CAGGACAACGCCCACACA (WT reverse), ACACTTGCCTCTACCGGTTC (mutant forward) and CTGAACTTGTGGCCGTTTAC (mutant reverse). *Ai9*;*R26 ^LSL-TdTomato^* were genotyped using the following primers: CTGTTCCTGTACGGCATGG (R26-tdTomato forward), GGCATTAAAGCAGCGTATCC (R26-tdTomato reverse), AAGGGAGCTGCAGTGGAGTA (Rosa WT forward), and CCGAAAATCTGTGGGAAGTC (Rosa WT reverse). Bax mice were genotyped using the following primers: GTTGACCAGAGTGGCGTAGG (Jax-Bax common reverse), CCGCTTCCATTGCTCAGCGG (Jax-Bax KO forward), and GAGCTGATCAGAACCATCATG (Jax-Bax WT forward).

### Animal and Tissue Procedures

#### Retinal Dissections

Mouse eyes were enucleated and drop fixed in 4% electron microscopy (EM) grade paraformaldehyde (PFA) in 1X phosphate buffered saline (PBS) for 5 minutes before the cornea was pierced with a 30G, 0.5 inch needle and placed back in PFA for an additional 25 minutes at 4°C. After fixation, the cornea was removed using microdissection scissors. For retinal cross-sections, retinal cups with the lens intact were washed three times for 5 minutes in 1X PBS with 0.1% sodium azide and cryoprotected in a 15% sucrose in 1x PBS solution overnight at 4°C. The next day, the lens was removed and retinal cups were placed in silicone embedding molds (SPI Supplies) filled with Neg-50 and flash-frozen with 2-methylbutane. Frozen retinal blocks were cut at 20 μm and sections were mounted on Fisher Superfrost slides using a Leica cryostat (CM3050S). For retinal flatmounts, after washing with three times for 5 minutes in 1x PBS, the cornea was cut with microdissection scissors and the sclera was removed, leaving only the retina and lens. The retina was then fixed for another 30 minutes in PFA before washing three times for 5 minutes in 1x PBS with 0.1% sodium azide before proceeding with flatmount immunohistochemistry.

For embryonic cross-sections, pregnant mice were euthanized by inhalation of CO_2_ followed by cervical dislocation. Embryos were extracted and immediately decapitated. Embryo heads were drop fixed in 4% EM-grade PFA in 1X PBS overnight at 4°C. On Day 2, tissue was washed three times for 15 minutes in 1x PBS and then placed in a 7.5% sucrose solution for 2 hours, followed by overnight in a 15% sucrose solution. On Day 3, tissue was placed in Peel-a-way embedding molds filled with Neg-50 and flash-frozen in 2-methylbutane. Tissue blocks were sectioned at 20 μm onto slides using a Leica cryostat (CM3050S).

### Immunohistochemistry

Slide edges were outlined in a hydrophobic ImmEdge pen. Slides were washed in 1x PBS for 5 minutes to remove the residual Neg-50 and incubated for 30 minutes in 1ml of blocking solution (5% Normal Donkey Serum, 0.2% Triton-X 100 in 1x PBS). Retinas were stained with primary antibody in blocking solution overnight at 4°C. Slides were then washed with 1x PBS three times for 10 minutes before applying secondary antibody in blocking solution for 2 hours at room temperature. Slides were then washed with 1x PBS two times, incubated in Hoechst in 1x PBS (1:5000) for 5 minutes and followed with a final 1x PBS wash, followed by application of Flouromount-G, coverslipping, and sealing the edges with clear nail polish.

For retinal flatmounts, retinal cups with the lens intact were initially incubated in blocking solution for 1 hour before incubating in primary antibodies for 5 days at 4°C. Retinas were then washed in 1x PBS three times for 1 hour before incubating in secondary antibodies overnight at 4°C, followed by three washes in 1x PBS for 1 hour each. The lens was removed from the retina cups, and the retinas were cut using an angled dissection knife into a cloverleaf shape to flatten the retina. Glass slides were covered with tape (VWR® Wafer Box Sealing Tape, Polyethylene – Transparent - #11211-934) featuring a square opening in the center to prevent tissue compression by the coverslip.

### RNAscope

Eyes were enucleated and immediately dropped in 4% EM grade PFA for 2 hours at 4°C, with corneas removed halfway through fixation. Afterwards, retinas underwent a sucrose gradient (5%, 10% and 15% in 1x PBS) each for 10-15 minutes on a rocker at room temperature before freezing in Neg-50 and 2-methylbutane. Retina blocks were cryosectioned at 20μm onto slides and left to fully dry before dehydrating in an ethanol gradient (25%, 50%, 75%, and 100%) for 5 minutes each in 1x PBS. Slides were immersed one final time in a slide mailer with fresh 100% ethanol for storage at - 20°C until use. Slides were thawed to room temperature and the RNAscope assay was conducted using the RNAscope^TM^ Multiplex Fluorescent Detection Reagents v2 kit according to the manufacturer’s protocol and the following modification: 1) Protease III was used for 20 minutes at room temperature.

### Fluorescence Image Acquisition

Retinal and embryonic cross-sections were imaged on a Zeiss Axio Imager M2 upright microscope equipped with an Apotome.2 and a metal halide light source (HXP 200 C), Axiocam 506 mono camera, and a 20X/0.8 NA Plan-Apochromat objective. For retinal cross-sections, images from the central retina were captured. Z-stack images were acquired in 0.5μm sections totaling 14-20μm thick. Retinal flatmounts were imaged using a 20x objective on a Zeiss LSM 980 laser scanning confocal built around a motorized Zeiss Axio Observer Z1 inverted microscope in the OHSU Advanced Light Imaging Core. Figures were composed in Affinity Designer v. 1.10.6 (now branded as Affinity by Canva).

### Quantification and statistical analysis Image Quantification

Littermate controls were used for all analyses, with the exception of experiments involving either triple-transgenic mice (*BAX* and *Hb9* mice) or *ChAT^Cre^*;*Isl1^cKO^* mice. Since the expected Mendelian frequency of each triple-transgenic genotype is low, control and mutant animals were rarely obtained within the same litter; these animals were therefore pooled across litters and age-matched. *ChAT^Cre^*;*Isl1^cKO^* mutant lethality limited the number of surviving mutants per litter and were also pooled.

#### Flatmount Cell Counts

Three regions of interest (ROIs) were selected from the central retina. For image processing, background subtraction (rolling ball = 50), despeckle and a median filter (radius = 2) were applied. The ROI size is indicated in figure legends. GCL and INL images were quantified from the same regions of the retina, differing in tissue z-depth only. Cell counts from single channel images (SOX2^+^ SACs, HCs) were counted by making z-stack images of the GCL and INL, followed by setting the ROI, and using the Multi-Point tool in Fiji/ImageJ.

#### IPLaminator Analysis

All fluorescence quantification of the IPL was assessed using IPLaminator, an ImageJ plugin designed to bin and quantify fluorescence intensity profiles specifically in retinal cross-sections (S. Li et al., 2016). For all quantification, Hoechst was used to define the IPL area whereas the fluorescent marker (e.g. ChAT, VAChT, CALB1, CALB2, etc) was measured. Using IPLaminator, the IPL was divided into 20 equal bins to normalize any variance in IPL thickness. For especially noisy images with high background (e.g. CALB1, GFP), three background ROIs were averaged to give an average background measurement that was then subtracted from each corresponding averaged technical replicate.

#### Colocalization Image Analysis

To assess colocalization across multiple markers, ROIs were first defined on either the SOX2 or tdT channel by applying an intensity threshold in Fiji/ImageJ to generate binary masks. These ROIs were then overlaid onto the remaining channel(s) of interest and cells were scored as double- or triple-positive if the remaining channel(s) of interest showed signal in the ROI. To assess differences in anti-ChAT fluorescence between *ChAT^Cre^;Isl1^Ctrl^*and *ChAT^Cre^;Isl1^cKO^* SACs, ROIs were made from 50 tdT^+^ cell bodies per mouse and overlaid onto the ChAT channel. Average integrated density measurements from Fiji/ImageJ were calculated and used for statistical analysis.

### Statistics

For all datasets, the variance was reported as mean ± S.E.M. The numbers of mice used for each analysis are referred to as “N” in the text or the figure legends. As the *Ptf1a^Cre^;Isl1^cKO^* phenotype (no SACs in the GCL) was readily apparent, blind analyses on SACs was not feasible. Other analyses on HCs, other ACs, BCs, and RGCs were performed blinded to genotype as there were no identifying differences between genotypes.

For comparisons between two groups, significance was determined using a two-tailed Students t-test. Comparisons between more than two groups were determined using a 2-way ANOVA with a Šidák post-hoc analysis. Statistical significance was set at α = 0.05 (p < 0.05). All statistical analyses and data visualization were performed with Microsoft Excel and GraphPad Prism (version 10.6.1).

### Key resources table

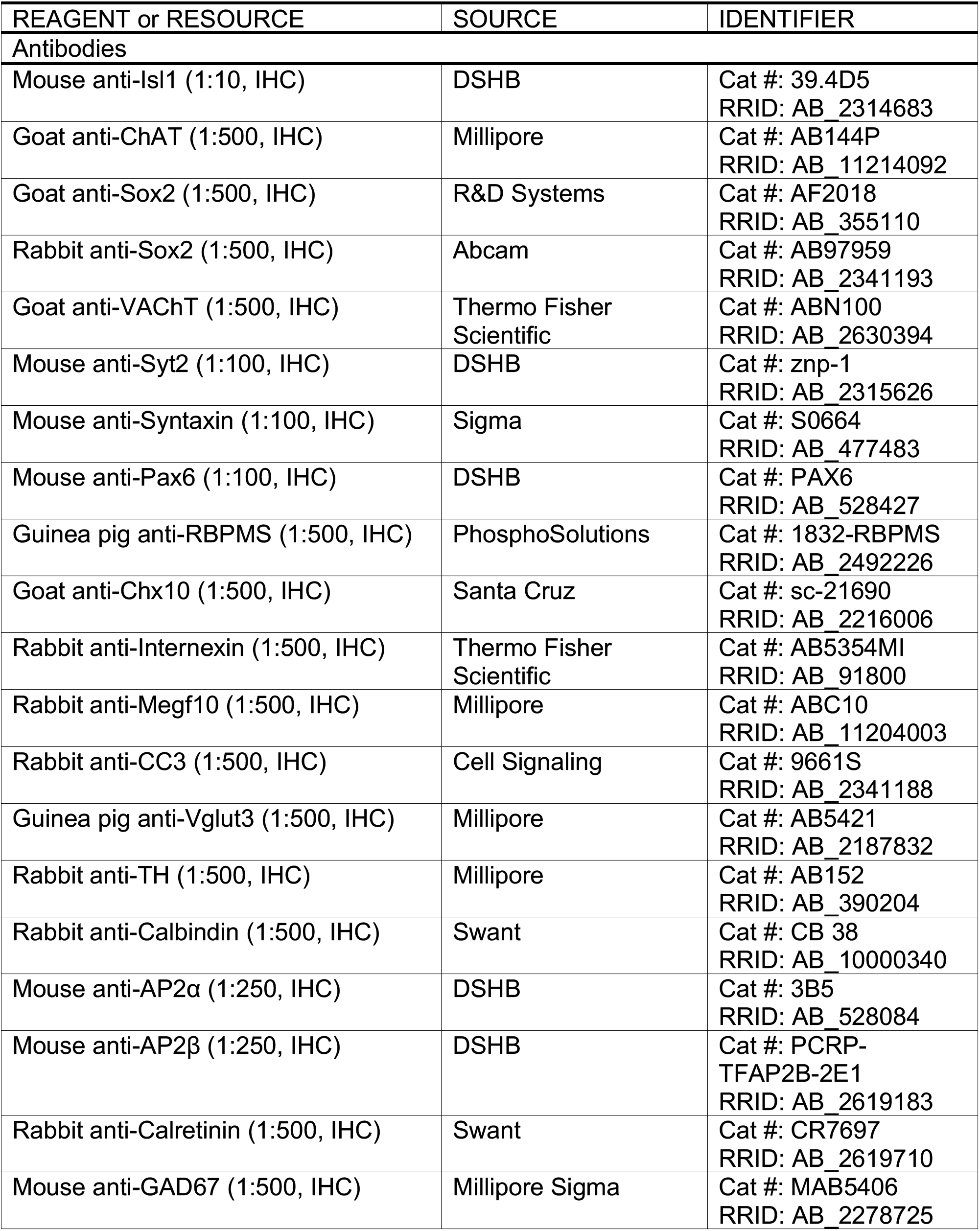
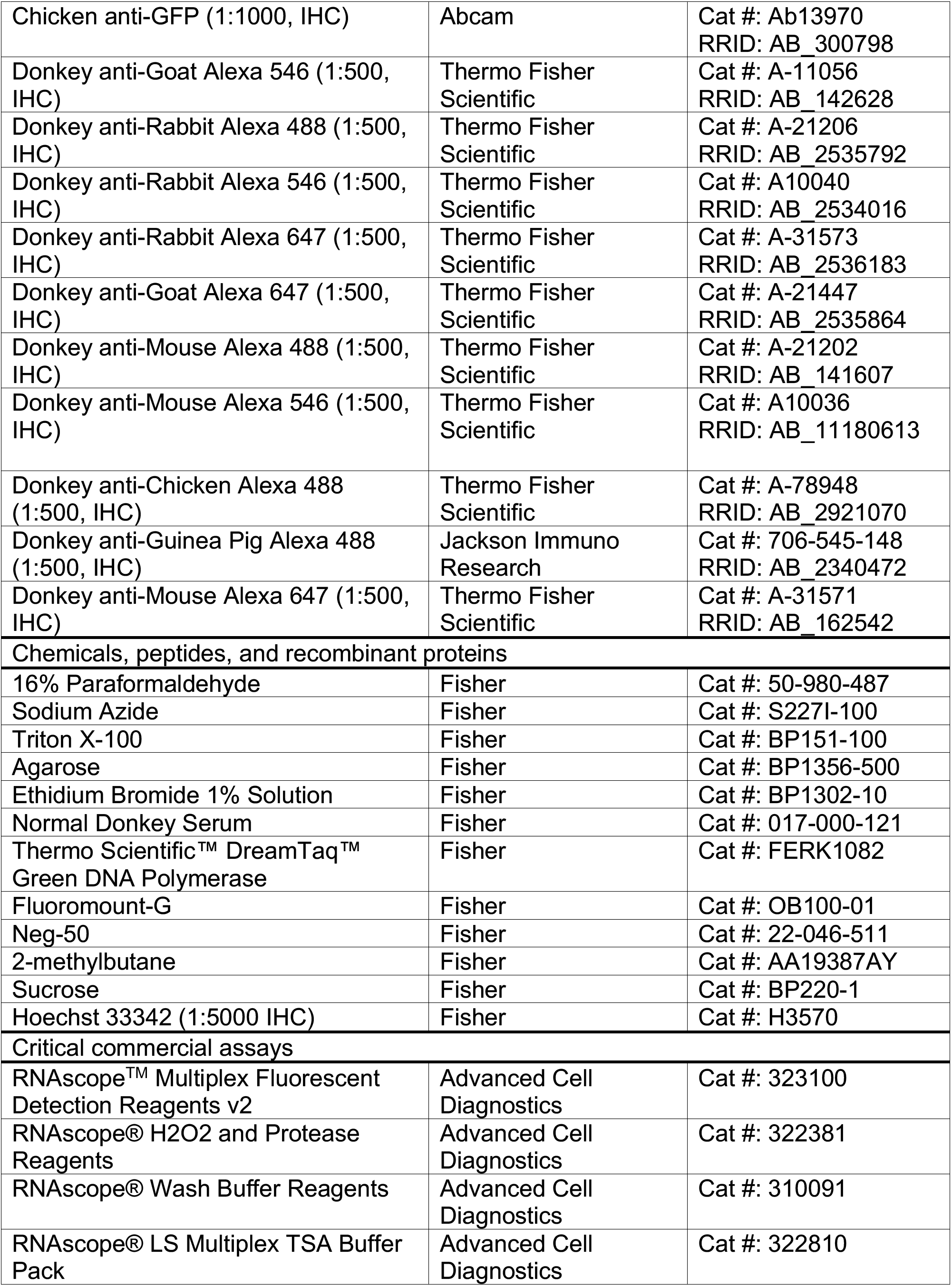
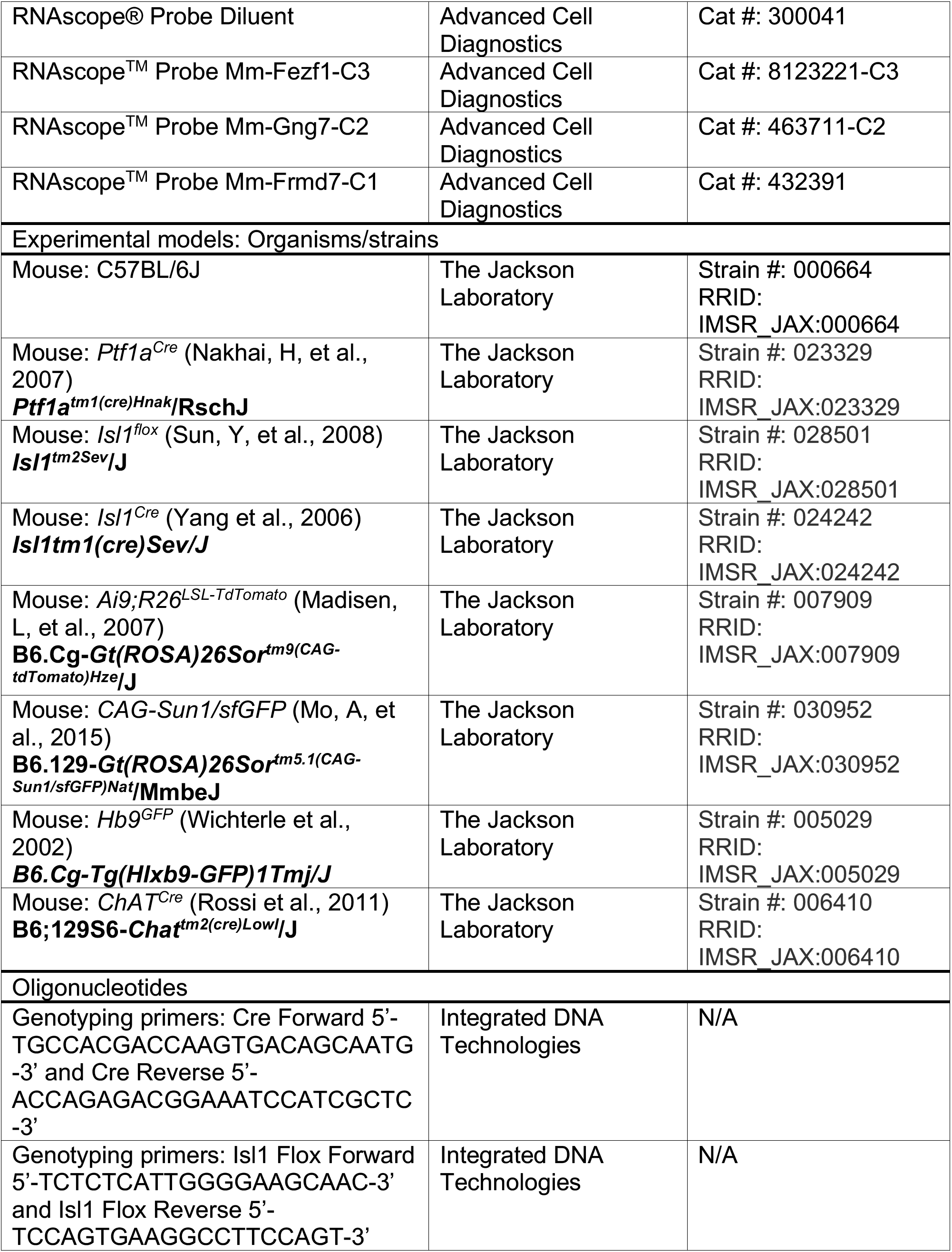
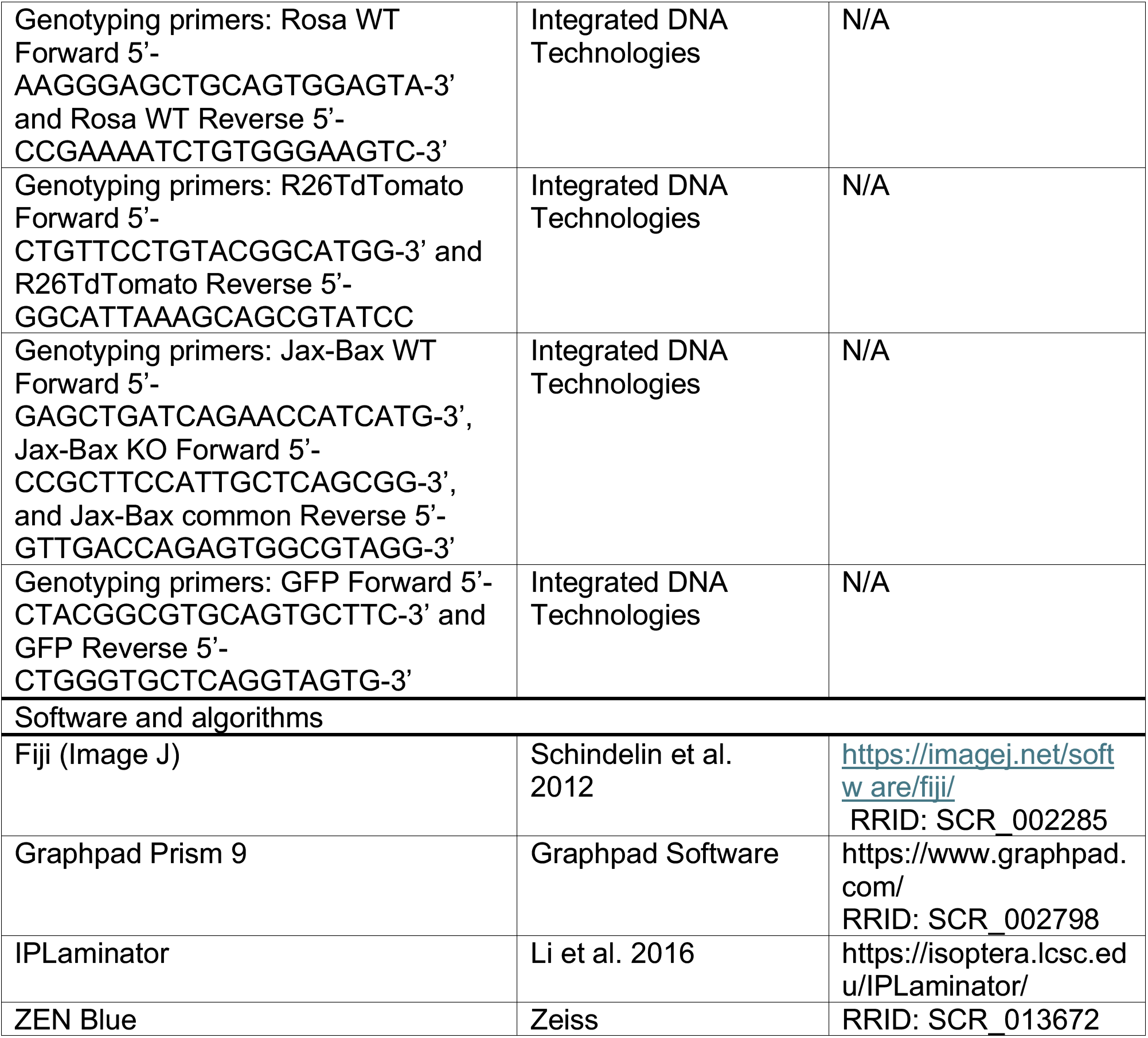

## Acknowledgements

This work was funded by NIH R01EY032057 (KMW), T32NS007466 (YSA), Lacroute Fellowship (YSA), HHMI Gilliam Fellowship (YSA).

---

**Supplementary Figure S1.**
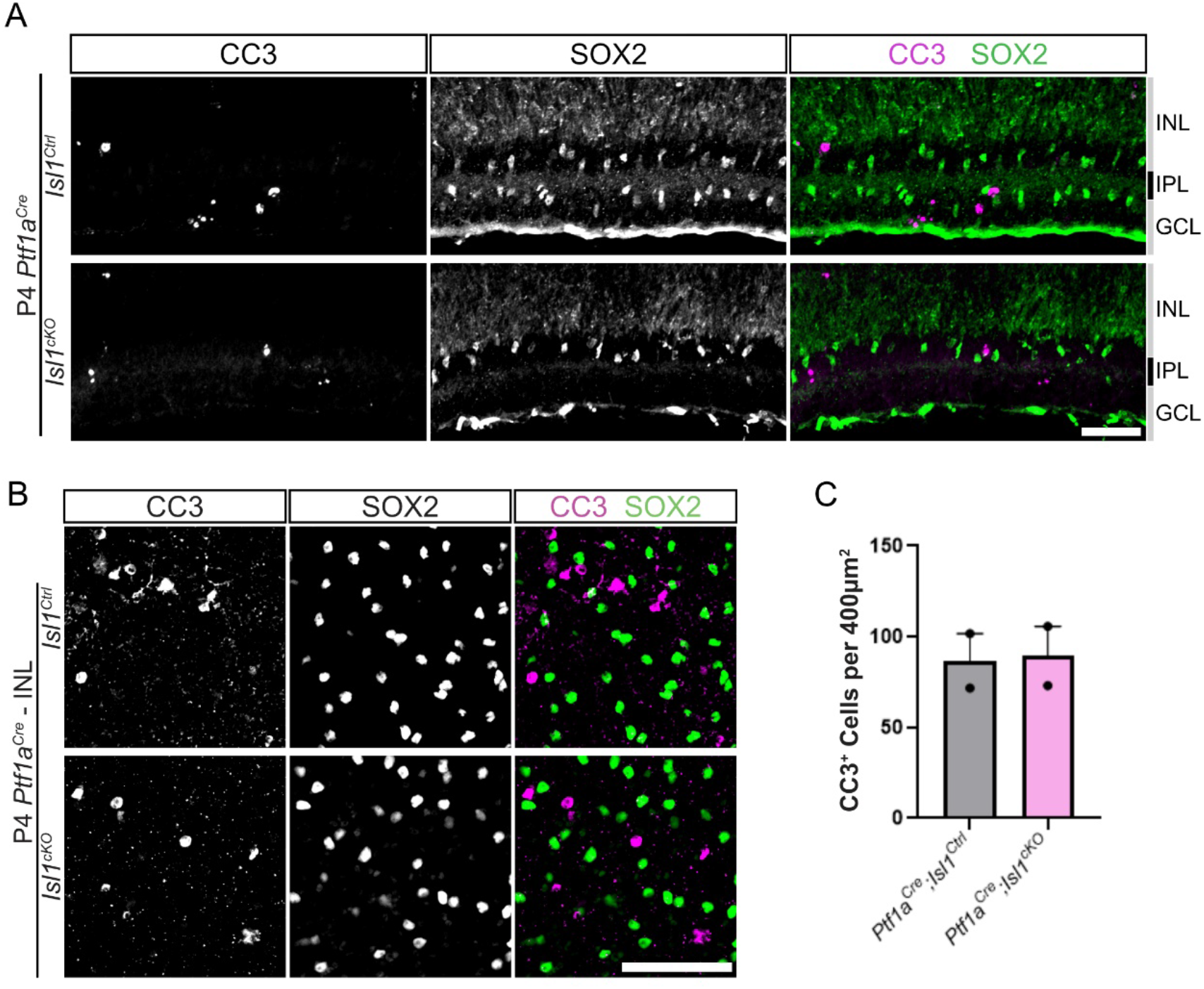
SACs are the only AC to express *Isl1*. (**A**) P28 *Isl1^Cre^;Sun1^GFP/+^* flatmount retinas immunostained for SAC-specific ChAT and GFP in the INL and GCL. Yellow arrows indicate GFP^+^ SACs in the GCL amongst GFP^+^ RGCs (n = 4). (**B-D**) P28 *Isl1^Cre^;Sun1^GFP/+^*cross sectioned retinas co-immunolabeled with (**B**) RBPMS (n = 4); (**C**) CHX10 (n = 4); (**D**) GFP, pan-AC AP2, and SOX2 (n = 4) scale bar = 50 μm. **(E)** P28 *Isl1^Cre^;Sun1^GFP/+^*flatmount retinas immunostained for AP2 and SOX2 in the INL and GCL. Yellow arrows indicate SACs; blue arrows indicate retinal astrocyte (n = 3); scale bar = 40 μm.

**Supplementary Figure S2.**
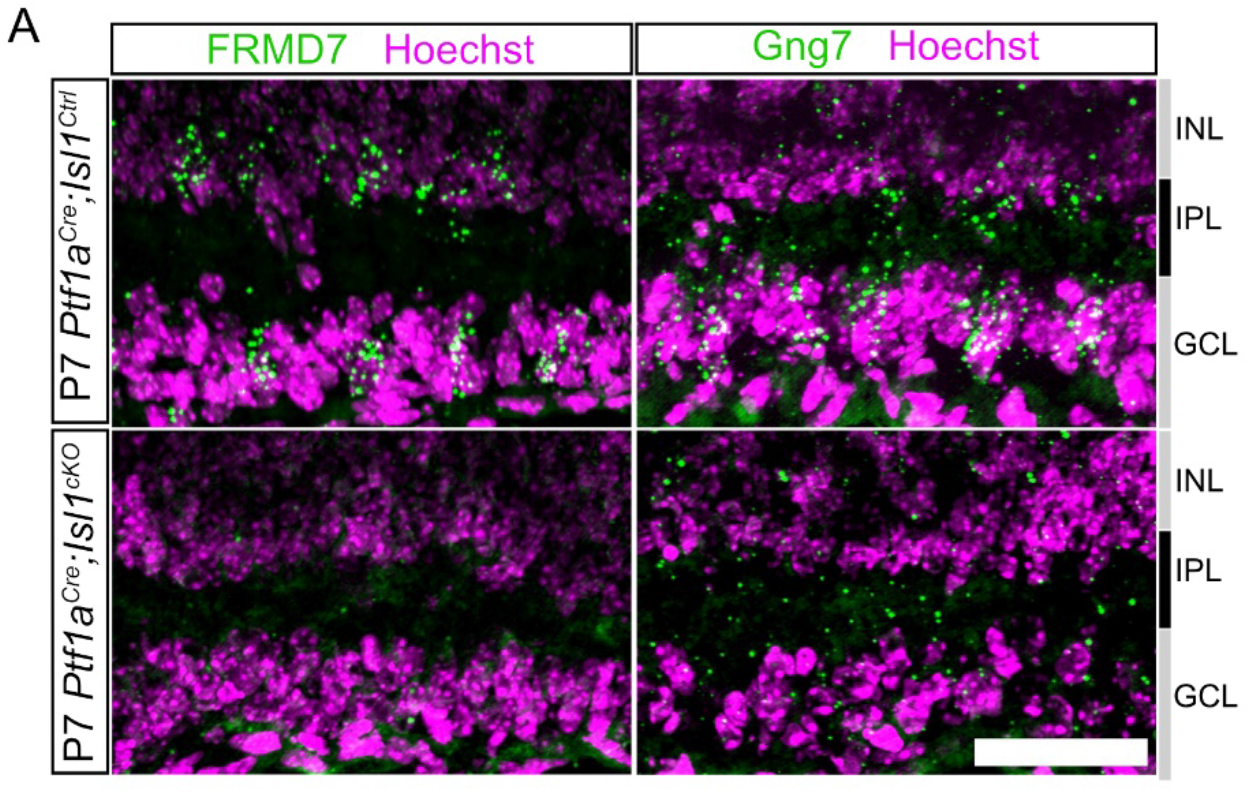
SACs are not CC3 positive in *Ptf1a^Cre^;Isl1^cKO^*retinas. Cross-sections (**A**) and INL flatmounts (**B**) of P4 *Ptf1a^Cre^;Isl1^Ctrl^*(top, n = 2) and *Ptf1a^Cre^;Isl1^cKO^* (bottom, n = 2) retinas immunostained for CC3 (magenta) and SOX2 (green). Scale bar = 50µm. (**C**) Quantification of CC3^+^ cell bodies in flamounts from *Isl1* control and mutant retinas.

**Supplementary Figure S3.**
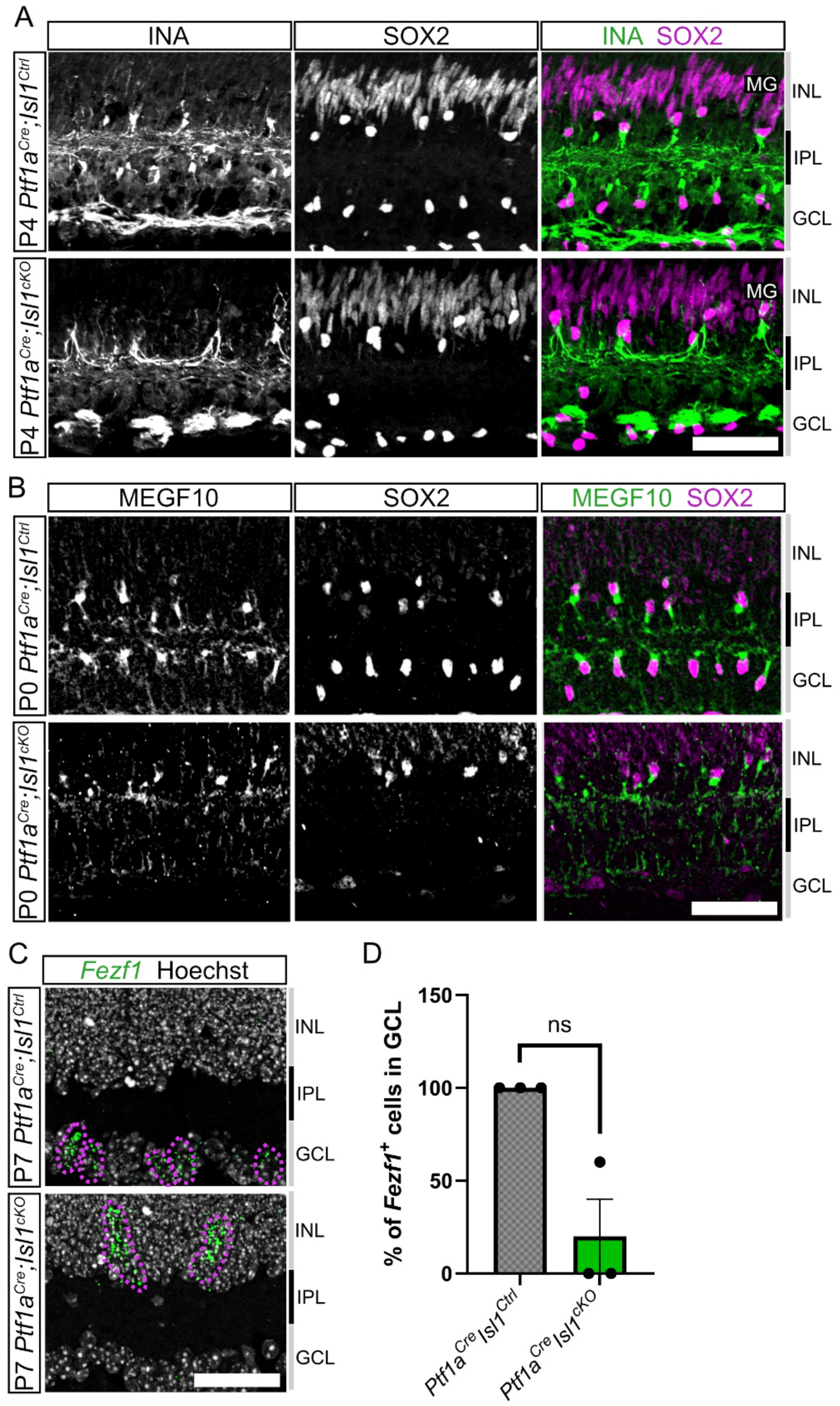
Expression of SAC-specific markers is eliminated in *Ptf1a^Cre^;Isl1^cKO^* retinas. (**A**) Fluorescence *in situ* hybridization (FISH) using probes against *FRMD7* or *Gng7* (green) in P7 *Ptf1a^Cre^;Isl1^Ctrl^* (top) (n = 3) and *Ptf1a^Cre^;Isl1^cKO^* (bottom) (n = 3) retinas. Scale bar = 50µm.

**Supplementary Figure S4.**
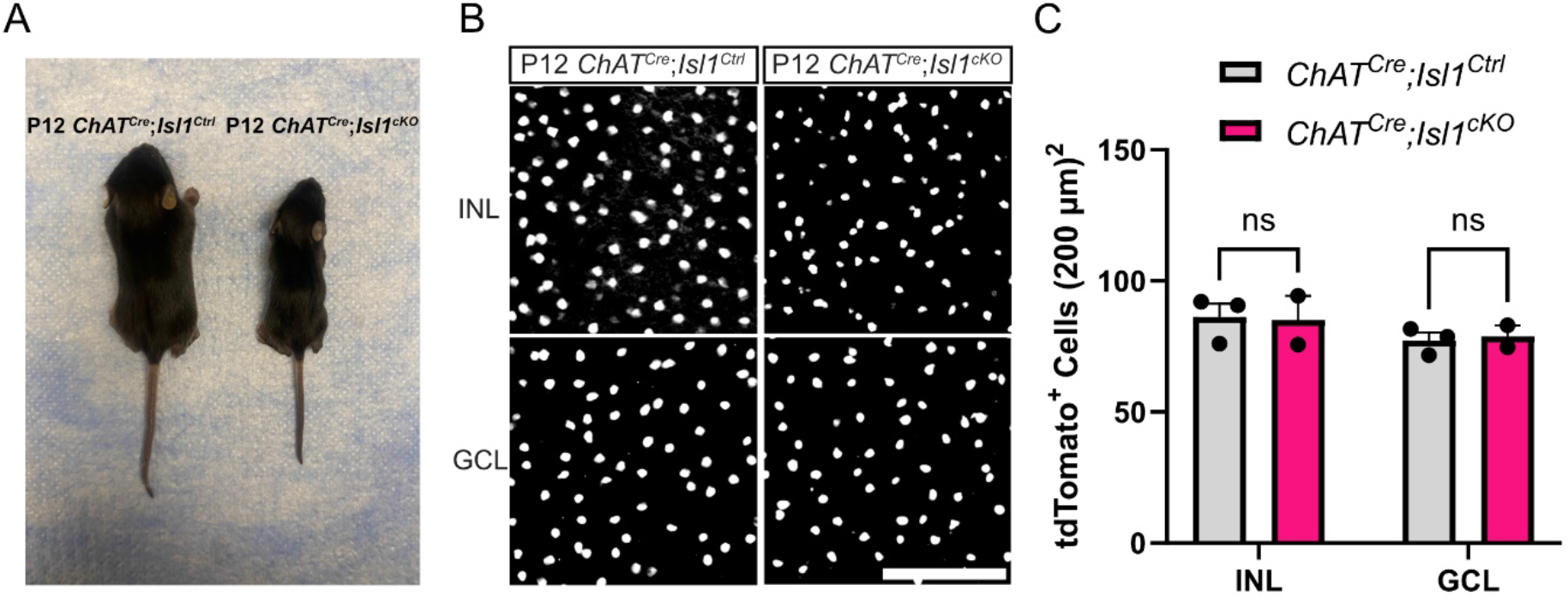
A subset of SAC markers are unaffected in *Ptf1a^Cre^;Isl1^cKO^* retinas. (**A**) Cross-sectioned P4 *Ptf1a^Cre^;Isl1^Ctrl^* (top) (n = 3) and *Ptf1a^Cre^;Isl1^cKO^* (bottom) (n = 3) retinas immunolabeled for INA (green) and SOX2 (magenta). (**B**) Cross-sectioned P0 *Ptf1a^Cre^;Isl1^Ctrl^*(top) (n = 1) and *Ptf1a^Cre^;Isl1^cKO^* (bottom) (n = 1) retinas immunolabeled for MEGF10 (magenta) and SOX2 (green). (**C**) Cross-sectioned RNAscope P7 *Ptf1a^Cre^;Isl1^Ctrl^* (top) (n = 3) and *Ptf1a^Cre^;Isl1^cKO^*(bottom) (n = 3) retinas for *Fezf1* (magenta) and Sox2 (green). Blue arrows = Fezf1^-^ OFF-SACs. Yellow arrows = Fezf1^+^ ON-SACs. Scale bar = 50µm. (**D**) Quantification of *Fezf1*^+^ cells in the GCL of P7 *Ptf1a^Cre^*;*Isl1^Ctrl^* (n = 3) and *Ptf1a^Cre^*;*Isl1^cKO^* (n = 3) retinas, expressed as a percentage of the control mean. Bars show mean ± SEM. One-sample t-test (p = 0.057).

**Figure S5.**
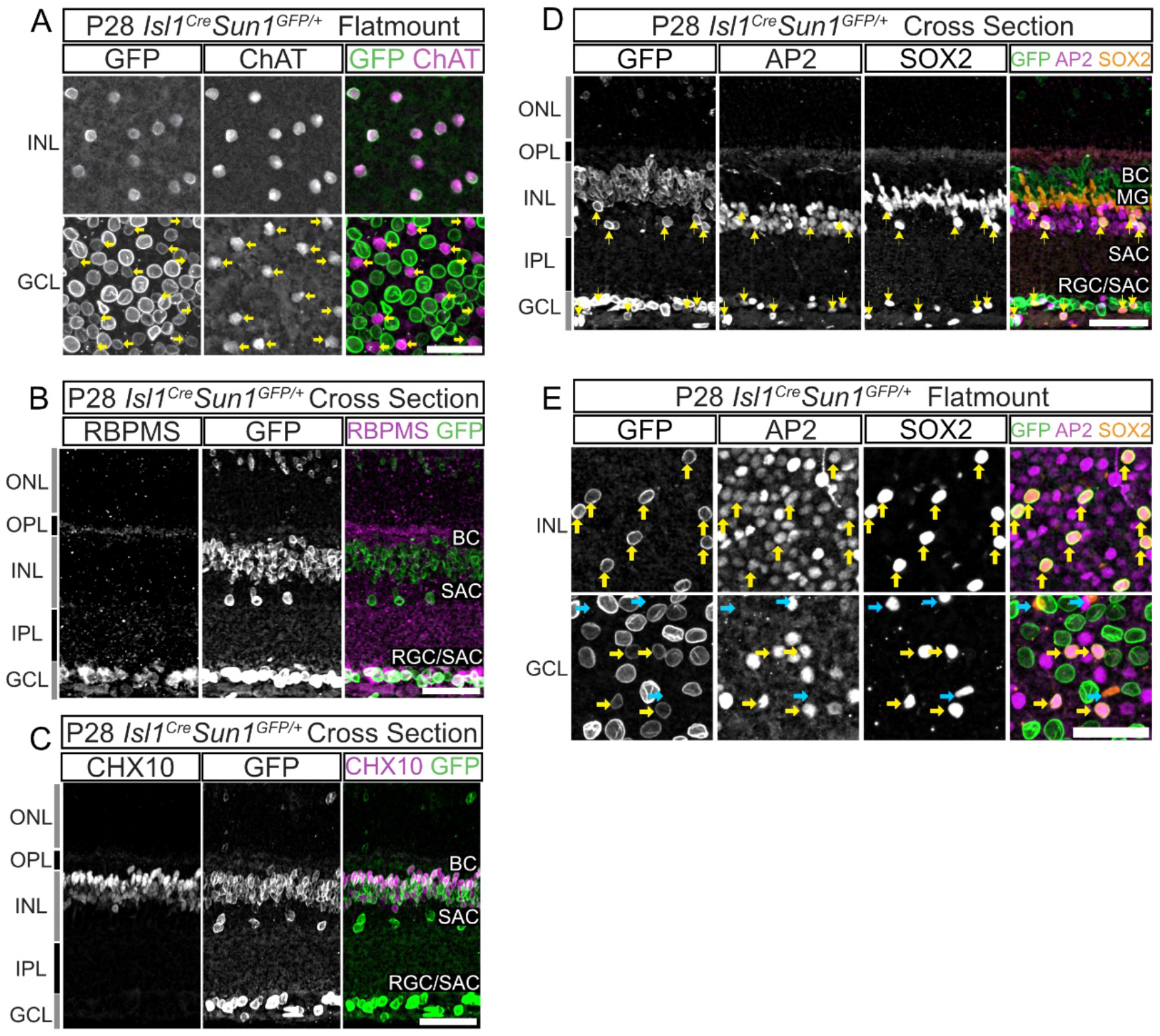
SAC population numbers are unaffected in post-developmental deletion of *Isl1*. (**A**) Image of P12 *ChAT^Cre(frt)^;Isl1^Ctrl^* and *ChAT^Cre(frt)^;Isl1^cKO^*littermates demonstrating size difference. (**B**) INL (top) and GCL (bottom) flatmount of endogenous tdTomato of *ChAT^Cre^*;*Isl1^Ctrl^* and *ChAT^Cre^*;*Isl1^cKO^*retinas. Scale bar = 100 μm. (**C**) Quantification of tdTomato^+^ SACs in the INL and GCL of P12 *ChAT^Cre^*;*Isl1^Ctrl^* (n = 3) and *ChAT^Cre^*;*Isl1^cKO^*(n = 2). Cells were counted in 200 x 200 μm fields. Genotypes were compared within each layer by two-way repeated-measures ANOVA (genotype x layer, with layer as the within-animal factor) followed by Šídák’s multiple comparisons test (INL: p = 0.9851; GCL: p = 0.9777).

**Supplementary Figure S6.**
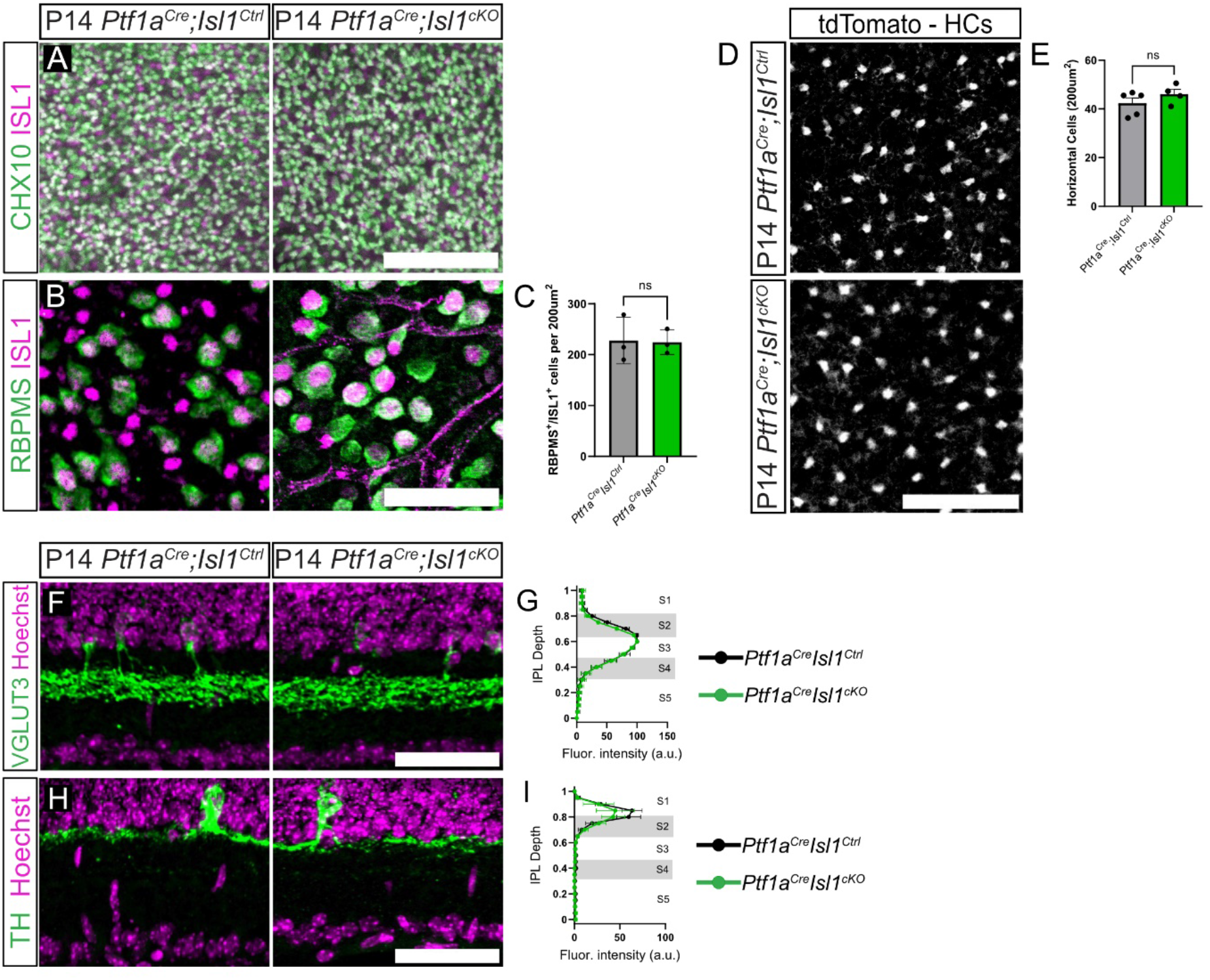
Other retinal populations are not affected *Ptf1a^Cre^;Isl1^cKO^* retinas. (**A**) Flatmounted P14 *Ptf1a^Cre^;Isl1^Ctrl^* (left) and *Ptf1a^Cre^;Isl1^cKO^* (right) retinas immunolabeled for pan-BC marker CHX10 (green) and ISL1 (magenta). Scale bar = 75µm. (**B**) Flatmounted P14 *Ptf1a^Cre^;Isl1^Ctrl^* (left) and *Ptf1a^Cre^;Isl1^cKO^*(right) immunolabeled with pan-RGC RBPMS (green) and ISL1 (magenta). Scale bar = 50µm. (**C**) Quantification of RPBMS^+^/ISL1^+^ cells between control (n = 3) and mutants (n = 3). (**D**) Flatmounted retinas showing tdTomato in the HC layer in P14 *Ptf1a^Cre^;Isl1^Ctrl^* and *Ptf1a^Cre^;Isl1^cKO^*retinas. Scale bar = 50µm. (**E**) Quantification of HC cell counts across control (n = 5) and mutant (n = 4) retinas. Cross sections of P14 *Ptf1a^Cre^;Isl1^Ctrl^*(left) and *Ptf1a^Cre^;Isl1^cKO^* (right) immunolabeled for (**F**) VGLUT3 (green) and (**G**) TH (green) and Hoechst (magenta). Mean intensity (SEM) of fluorescence of VGLUT3^+^ dendritic lamination (**F**, controls, n = 3; mutants, n = 7) and TH^+^ dendritic lamination (**G**, controls, n = 6; mutants, n = 5) in the IPL. Scale bar = 50µm.

